# Optogenetic manipulation of individual or whole population *Caenorhabditis elegans* worms with an under hundred-dollar tool: the OptoArm

**DOI:** 10.1101/2021.03.19.435933

**Authors:** M. Koopman, L. Janssen, E.A.A. Nollen

**Affiliations:** European Research Institute for the Biology of Ageing, University of Groningen, University Medical Centre Groningen, The Netherlands

**Keywords:** OptoArm, Optogenetics, *Caenorhabditis elegans*, neuronal ageing, rhodopsin, worm-trackers

## Abstract

Optogenetic tools have revolutionized the study of neuronal circuits in *Caenorhabditis elegans.* The expression of light-sensitive ion channels or pumps under specific promotors allows researchers to modify the behavior of excitable cells. Several optogenetic systems have been developed to spatially and temporally photoactivate light-sensitive actuators in *C. elegans*. Nevertheless, their high costs and low flexibility have limited wide access to optogenetics. Here, we developed an inexpensive, easy-to-build, and adjustable optogenetics device for use on different microscopes and worm trackers, called the OptoArm. The OptoArm allows for single- and multiple-worm illumination and is adaptable in terms of light intensity, lighting profiles and light-color. We demonstrate the OptoArm’s power in a population-based study on contributions of motor circuit cells to age-related motility decline. We find that functional decline of cholinergic neurons mirrors motor decline, while GABAergic neurons and muscle cells are relatively age-resilient, suggesting that rate-limiting cells exist and determine neuronal circuit aging.

## Introduction

Neurotransmission is defined as the process by which neurons transfer information via chemical signals at synaptic contacts (e.g. synapses) with target cells. Those target cells can be other neurons, but also non-neuronal cell types (e.g. muscle cells). *Caenorhabditis elegans* has proven to be an important model organism to study fundamental neurobiology, including neurotransmission at chemical synapses (Rand et al., 1984; 1985; Lewis et al., 1987; Hosono et al., 1989; 1991; Maruyama et al., 1991; Nonet et al., 1993, 1998; Jorgensen et al., 1995; Miller et al., 1996; Goodman et al., 1998, 2012; Richmond et al., 1999a, 1999b, 2006, 2009; Francis et al., 2003; 2006). As a matter of fact, important molecular players involved in synaptic transmission, synaptic vesicle docking, priming, fusion and recycling have actually been discovered in *C. elegans* and are highly conserved in mammalian systems (Brose et al., 1995; Betz et al., 1998; Aravamudan et al., 1999; Augustin et al., 1999; Richmond et al., 1999b; Lackner et al., 1999; Sassa et al., 1999). Initially, pharmacological assays were predominantly used to analyze synaptic transmission and its associated molecular substrates in *C. elegans* and to determine whether an observed neurotransmission defect was pre- or postsynaptic of nature (Lewis et al., 1987; Miller et al., 1996). Then, the development of electrophysiology resulted in new techniques that have allowed researchers to study the electrical events that occur at synapses more directly (Richmond et al., 1999a, 1999b, 2006, 2009; Francis et al., 2003; 2006; Mellem et al., 2002; O’Hagan et al, 2005; Ramot et al., 2008a; Kang et al., 2010; Geffeney et al., 2011; Kawano et al., 2011; Goodman et al., 2012). These methods provide a quantifiable readout for the underlying synaptic activity and a way to investigate synaptic properties. Nevertheless, standard electrophysiology in *C. elegans* is technically challenging and cannot be performed in intact worms, making it impossible to directly correlate synaptic activity with behavioral readouts (Husson et al., 2013). The development of optogenetics, however, addressed this issue and created the possibility to directly manipulate synaptic activity and simultaneously look at behavioral changes (Nagel et al., 2005; Liewald et al., 2008; Liu et al., 2009; Schultheis et al., 2011a; 2011b; Husson et al., 2013).

Optogenetics is based on the genetic expression of a light-sensitive actuator that can actively influence biochemical reactions or neuronal activity in response to a non-invasive light stimulus (Zhang et al., 2007; Husson et al., 2013). The most common used ‘actuators’ are the rhodopsins, which have been discovered in algae, and are now widely used in different cells and organisms to depolarize or hyperpolarize cells upon light stimulation (Nagel et al., 2003, 2005, Li et al., 2005; Schroll et al., 2006; Bi et al., 2006; Ishizuka et al., 2006; Zhang et al., 2007; Fang-Yen et al., 2015). The transparent body of *C. elegans*, its amenability for genetic manipulation and its invariant nervous system, make the worm ideal for optogenetic manipulation (Hope et al., 1999; Kaletta et al., 2006; Antoscheckin et al., 2007; Husson et al., 2013). In fact, *C. elegans* was the first multicellular organisms to have its behavior manipulated *in vivo* by the photoactivation of channelrhodopsin-2 (ChR2) in muscle cells (Nagel et al., 2005). ChR2 is a light-sensitive cation channel that can undergo a light-induced conformational change, thereby allowing H^+^, Na^+^, K^+^ and Ca^2+^ ions to passively diffuse down their concentration gradients (Figure 1A) (Nagel et al., 2003, 2005; Kolbe et al., 2000). In excitable cells, this results into rapid depolarization of the plasma membrane and the subsequent initiation of downstream events, like fibril contraction in muscle cells and synaptic vesicle release in neuronal cells (Nagel et al., 2005; Zhang et al., 2007; Liewald et al., 2008; Almedom et al., 2009; Liu et al., 2009; Gao et al., 2011; Schultheis et al., 2011b; Kittelman et al., 2013). The specificity of the response is acquired by expressing optogenetic proteins under specific promoters (Figure 1B) in the worm. For instance, the expression of ChR2 in motor neurons (*unc-47, unc-17*) or muscles (*myo-3*) elicits either a flaccid or spastic paralysis upon complete illumination of the worms (Zhang et al., 2007; Liewald et al., 2008; Husson et al., 2013). This obvious and robust response results in a change in body length that can be used as a clear readout (Figure 1C) (Zhang et al., 2007; Liewald et al., 2008). Multiple other strains exist, in which for example specific interneurons or sensory neurons are under the control of ChR2 (Nagel et al., 2005; Franks et al., 2009; Guo et al., 2009; Narayan et al., 2011; Lindsay et al., 2011; Milward et al., 2011; Piggott et al., 2011; Leifer et al., 2011; Stirman et al., 2011; Faumont et al., 2011; Husson et al., 2012a; Busch et al., 2012; Schmitt et al., 2012). Functional ChR2 requires the chromophore all-trans retinal (ATR), which is not endogenously produced by *C. elegans*. Exogenous feeding of ATR can circumvent this problem and ensures functional ChR2 (Nagel et al., 2005)

**Figure 1:**
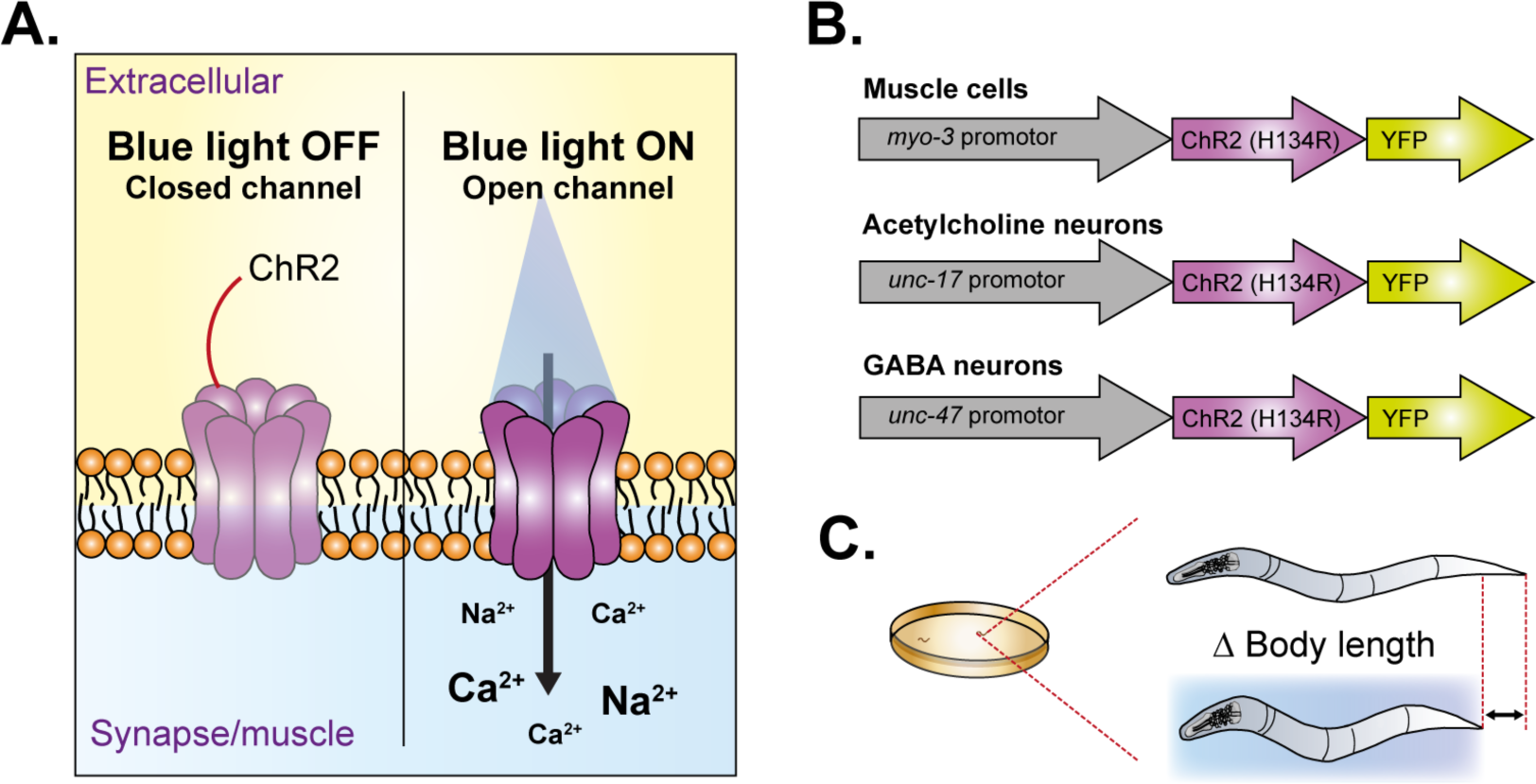
Channelrhodopsin-2 is a light-sensitive cation channel. **A)** Channelrhodopsin-2 (ChR2) is rapidly opened by stimulation with blue (440-460 nm) light in an essentially nondesensitizing manner. ChR2 is permeable by multiple cations like Na^+^ and Ca^2+^ and enables strong and rapid membrane depolarization of the cell it is expressed in. **B)** Multiple ChR2-expressing C. elegans strains exist, including but not limited to expressing in muscle cells (*myo-3*), cholinergic neurons (*unc-17*) and GABAergic neurons (*unc-47*). **C)** The change in body length is often used as a read-out of ChR2 activation (see B).

Over the years, multiple systems have been developed to illuminate *C. elegans* in a temporal and/or spatial manner for optogenetic experiments. Most of those systems are custom built and include fluorescence microscopes, or systems with lasers and shutters (Leifer et al., 2011; Weissenberger et al., 2011; Stirman et al., 2010; 2011; Husson et al., 2012b; Qiu et al., 2015; Rabinowitch et al., 2016a; Gengyo-Ando et al., 2017). These systems require costly equipment and are mostly designed for single-animal illumination. While the costs might limit the general use of these systems for a general audience, they do, however, provide clear benefits for fundamental research requiring single-cell stimulation. For example, some of these systems provide ways of following single worms in space and time with chromatic precision, thereby ensuring exact illumination of a specified anatomical position (Stirman et al., 2011; Leifer et al., 2011; Kocabas et al., 2012; Gengyo-Ando et al., 2017). This targeted illumination makes it possible to optogenetically excite single neurons for which specific promotors are not known (Kocabas et al., 2012). Moreover, in these expensive systems, one can adjust the light intensity relatively easily, making the systems flexible and adaptable for different types (e.g., short- or long-term) of experiments.

Several researchers have tried to make optogenetical experiments more accessible to a wide audience by successfully developing inexpensive (<1500 dollars) optogenetic systems (Pulver et al., 2011; Kawazoe et al., 2013; Rabinowitch et al., 2016b; Pokala et al., 2018; Crawford et al., 2020). Some of these low-cost systems even allow multiple worms to be illuminated at the same time (Kawazoe et al., 2013; Crawford et al., 2020). When considering the current developments of automated worm trackers (Tsibidis et al., 2007; Ramot et al., 2008b; Swierczek et al., 2011; Stirman et al., 2011; Leifer et al., 2011; Wang et al., 2013; Kwon et al., 2013; Restif et al., 2014; Javer et al., 2018; Perni et al., 2018; Koopman et al., 2020), the ability to illuminate multiple worms at the same time during tracking offers a clear benefit for studying both population and single-worm characteristics. However, there are also clear limitations of these low-cost systems. In fact, most of the systems require manual adaptations (e.g., aluminum foil, closed boxes) to ensure a light intensity that is high enough (e.g. 1.6 mW/mm^2^, or at least 1.0 mW/mm^2^) for optogenetic purposes in *C. elegans* (Rabinowitch et al., 2016b; Crawford et al., 2020). These adaptations make the simultaneous tracking of illuminated worms challenging and highly reduce the adaptability of the system to other experimental set-ups. Moreover, within low-cost systems the light-source often has to be placed in very close proximity (<2 cm) to the worms, which makes such systems not suitable for long-term stimulation due to potential heat generation and phototoxicity (Kawazoe et al., 2013; Rabinowitch et al., 2016b; Crawford et al., 2019). Some systems, such as the OptoGenBox (Busack et al., 2020), are exempt from these issues, but they are also more expensive.

Clearly, there is no shortage of tools and techniques to perform optogenetic experiments with *C. elegans*. Ideally, one would simply look at all the advantages and limitations of each system and select for the platform that fits best to the biological questions to be answered. Practically, however, there are often different types of biological questions to be answered or various approaches required to tackle a specific hypothesis. We argue that the use of optogenetics should not be restricted by limited capabilities of single system or available (financial) resources. From that perspective there remains a challenge in developing low-cost optogenetic devises with high flexibility and adaptability. Here, we describe in detail how we developed an under hundred-dollar optogenetic device, the OptoArm. This system provides the user with an easy-to-build, highly adaptable, flexible and reliable optogenetic system. The arm allows integration in different experimental set-ups and tackles both single- and multiple-worm illumination, making combinatory approaches and population-based studies possible. In fact, we demonstrate the use of the OptoArm by determining the functional contribution of motor circuit cells to the age-related decline in motility. Finally, by offering both the option to build a manual and a fully automated system, researchers are able to pick the set-up that fits their budget and requirements best. The biologically and technically validated OptoArm makes optogenetics experiments accessible for researchers both within and outside the laboratory, including teaching institutions.

## Results

### Efficient thermal management is key in constructing a compact, inexpensive set-up

#### Construction

To build a simple and inexpensive optogenetics device that can be adjusted to laboratory-specific parameters, specific applications and requirements, we investigated the use of low-cost, high-intensity LEDs with interchangeable optics. The ease of the construction and the low costs of the system ensures accessibility for both researchers and students. We opted for a straightforward electronic circuitry (Figure 2A) using a 700 mA LED-driver to ensure the feeding of fixed electric current to a 1.03 W LED with a wavelength ranging between 440-460 nm (optimal: 448 nm). The placement of the ON/OFF switch between the CTRL and REF pins, in parallel with the potentiometer (Figure 2A, main panel), provides the best management of inrush power and spikes to the LED. Therefore, this is the recommended configuration to ensure an optimal lifetime for the LED. For prototyping purposes, however, we used a circuit with the switch serially connected to the LED driver and the DC plug (Figure 2A, insert). As depicted in Figure 2B, in addition to the high-intensity LED and the LED-driver, only a small set of components is required (see Table 1 and *Material and methods*): a potentiometer to adjust intensity, an ON/OFF switch, a lens holder (home-made) with interchangeable lenses, a heatsink and thermal adhesives to connect the components (see *thermal stability* section) and a 9V_DC_ adapter to power the system. The different steps required to build the optogenetic arm can be found in Figure 2C and Table 2. Eventually, we installed the complete circuitry on a standard flask clip clamp, as this could be used in combination with different stands for different set-ups (figure 2D, E, Table 1). Note that the combination of the wavelength and high intensity makes the LED dangerous for the human eye. Try to avoid looking directly into the LED, use protective sheets or wear protective glasses.

**Figure 2:**
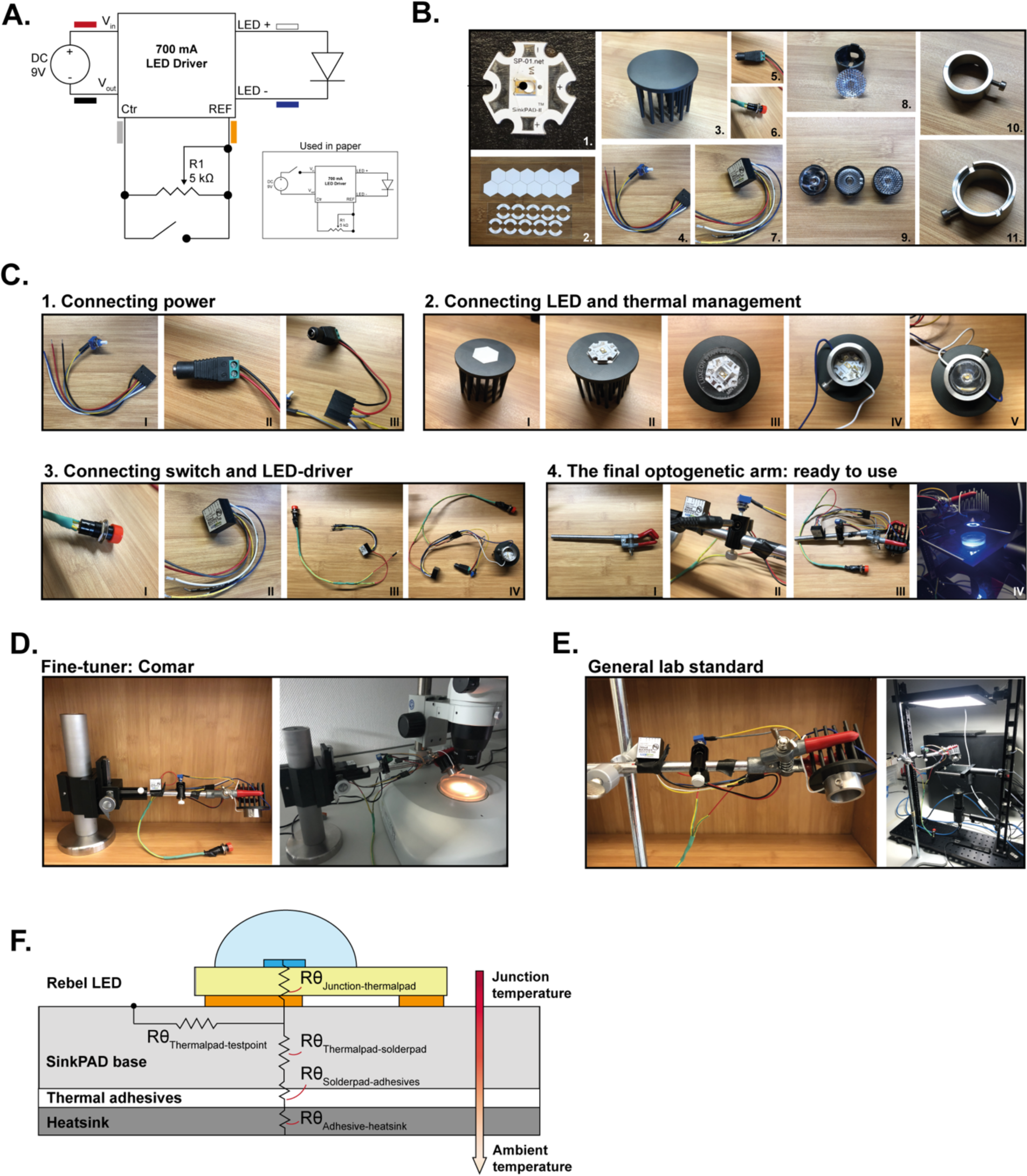
Building the OptoArm only requires basic knowledge of electronics and thermal management. **A)** The optimal electronic circuitry of the OptoArm. Inset: the variant of the circuitry used in this paper to test and validate the system. The color codes refer to the wires of the LED-driver that is used in this paper. **B)** The essential components required to build the electronic circuitry of the OptoArm: 1. Luxeon Rebel LED, 2. Thermal adhesives, 3. Heatsink, 4. Wire harness with potentiometer, 5. DC-plug, 6. ON/OFF button, 7. LED driver, 8. lens with case, 9. Several lenses, 10. and 11. Lens holder. **C)** The steps required to construct the OptoArm, detailed instructions per picture can be found in Table 2. *STEP 1:* connecting the solderless DC-plug to the connecting wire harness. *STEP 2*: mounting the LED on a heatsink and soldering the connecting wire harness to the LED cathode and anode. *STEP 3*: Connecting the ON/OF switch to the LED driver and connecting the driver to the wire harness. Note, the pictures show a serial set-up, and not the recommend parallel wiring (see 2A, main). *STEP 4*: mounting the electronic circuitry to a standard flask clamp to finish the OptoArm. By placing the OptoArm in either a **D)** fine-tuner or a **E)** general lab standard, the system can be used for different applications. **F)** The different thermal resistances (*Rθ*) in the heat flow between the LED junction, temperature test point and bottom of the LED assembly.

**TABLE 1:** Different variants of the OptoArm, their requirements and costs.

|  | Minimal requirements<br>OptoArm<sup>a</sup><br>(Figure 2) | Minimal requirements<br>OptoArm with low-cost<br>stand<sup>a,b</sup><br>(Figure 2) | Automated OptoArm<br>with low-cost stand<sup>a</sup><br>(Figure 10) | Additional improvements<br>or available changes<sup>a, d</sup> |
| --- | --- | --- | --- | --- |
| **Electronics** | >Royal-Blue (448nm) Rebel LED<br>>700 mA externally dimmable BuckPuck DC driver<br>>Connecting wire with adjustable potentiometer of 5 kΩ<br>>Latching pressure switch (ON/OFF switch)<br>>DC contra plug<br>>9V<sub>DC</sub> – Adapter | **All the previously mentioned components** | **All the previously mentioned components except:**<br>>Connecting wire with adjustable potentiometer of 5 kΩ<br>**Plus:**<br>> Arduino uno (or clone)<br>> Capacitor (0.1 uf)<br>> Adafruit LCD-shield | **Yellow LED for NpHR/Halo/Mac<sup>c</sup>:**<br>e.g Lime (567nm) Rebel LED on a SinkPAD-II 20mm Star Base - 368 lm @ 700mA<br>COST: \$11<br>**Tri-star LED for higher intensity<sup>c</sup>:**<br>Royal-Blue (448nm) Rebel LED on a SinkPAD-II 20mm Tri-Star Base - 3090 mW @ 700mA.<br>COSTS: \$14 |
| **Heat management** | >Heatsink 5.2 °C/W<br>>Pre-cut thermal adhesive tape | **All the previously mentioned components** | **All the previously mentioned components** | - |
| **Optics** | >Khatod 10° lens<br>>Fraen 21° lens<br>>Khatod 40° lens | **All the previously mentioned components** | **All the previously mentioned components** | - |
| **Mounting** | >Lens-holder<br>>Three-pronged clamp | **All the previously mentioned components AND:**<br>> Stand rod<br>> Double boss-head<br>> Retort stand base, tripod | **All the previously mentioned components** | **Comar fine-tuner stand<sup>f</sup>:**<br>Basic carrier, pinion stage, post-holder, rod, self-manufactured base. COSTS: ~ \$ 280<br>**Cross-clamp to fixate DC-plug on arm<sup>g</sup>:**<br>COSTS: ~ \$12 |
| **Average costs** | **~ \$ 65** | **~ \$ 90** | **~ \$ 128<sup>c</sup>**<br>(Arduino clone: \$ 116) | |

<sup>a</sup> For components originally listed in euros, we calculated the dollar prize with the current exchange rate in mind. All prices are excluding soldering material and standard materials such as extra wires and shrink tubing. Also, standard worm equipment is not taken into account (e.g. microscopes and worm trackers). <sup>b</sup> The low-cost stand provides a way to work with the WF-NTP and a microscope set-up and is therefore the most flexible. <sup>c</sup> The price is based on an original Arduino uno R3. Clones tend to be much cheaper, but identical in the board lay-out. <sup>d</sup> The components are not essential, but potential substitutes or additions. <sup>e</sup> When a Tri-Star LED or any other LED is used, we highly recommend to re-evaluate the thermal management. The heatsink of the OptoArm was specifically selected based on the specification of a single royal-blue LED. <sup>f</sup> The Comar fine-tuner stand is a more expensive stand, but allows fine-adjustments of height due to the pinion stage. All the specifications can be found in the method section. <sup>g</sup> We did use the cross-clamp to organize the OptoArm and fixate the DC contra plug, see Figure 2C.

**TABLE 2:** Instructions for building the OptoArm (see Figure 2C)

| Steps | Goal | Instructions<sup>a</sup> | Critical notes |
| --- | --- | --- | --- |
| Step 1 | Connecting power | In order to connect a DC-adaptor to the optogenetics system, we use a soldering-free DC-plug. The black and red wires of the connecting wire harness (I) should be attached to the DC-plug (II, III). Note that the used wire harness has already a 5 kΩ potentiometer pre-installed. When using a different system, make sure to manually add a potentiometer parallel to the switch between the yellow (REF) and grey (CTRL) wires. | Determine the center and barrel polarity of the system to ensure appropriate wiring (with a multimeter) |
| Step 2 | Connecting LED and thermal management | For appropriate thermal management use a heatsink that fulfils the criteria of the system (see *thermal management*, box 1). Clean the heatsink with ethanol before adding a pre-cut thermal adhesive (I). Remove the top-layer of the thermal adhesive and place the LED on top of it (II). Apply firm pressure for at least 30 seconds with an assembly press that's included with the single rebel led (III). Solder the blue and white wires of the connecting wire harness to the cathode and anode of the LED respectively. Then, cut some pre-cut thermal adhesive in the shape of the lens-holder and attach the lens-holder via the tape to the heatsink (IV). Finally, by loosening the screw of the lens-holder, lenses can be added to or removed from the system (V). | It is important to solder when the LED is already on the heatsink to ensure heat-dissipation and circumvent overheating of the LED |
| Step 3 | Connecting switch and LED driver | Add the ON/OFF switch (I) and LED driver (II) to the system (III, IV). The driver can be connected to the wire harness by soldering male pin connectors to the wires of the LED-driver and clicking them in the connecting wire harness. Note, that the pictures (Figure 2C) show the serial set-up used in the paper, in which the red-wire coming from the DC-plug is connected with the ON/OFF switch before connecting to the LED-driver. We recommend to place the switch in parallel to the potentiometer between the yellow and grey wires (See Figure 2A). | The placement of the ON/OFF switch between the CTRL and REF pins, in parallel with the potentiometer provides the best management of inrush power and spikes to the LED. |
| Step 4 | Mounting on the arm: the final OptoArm | Add the whole circuitry to a standard flask clamp (I). Use a cross clamp to fixate the DC-plug on the arm (II, III). By placing the OptoArm in either a fine-tuner or a general lab standard, the system can be used for different applications. | - |

<sup>a</sup>The different steps and roman numerals refer to those used in Figure 2C.

The resulting OptoArm is compact in nature, making it easy to move around and integrate within different recording systems. The prices in Table 1 are an approximation based on the main components. They do not include the required, basic soldering materials, extra wires, shrink tubing and pin-adaptors, nor standard *C. elegans* equipment, like a microscope. We were able to reduce our costs further (<65 dollar) by recycling available material in the lab, including the flask clip clamp and different stands. Therefore, it should be underlined that almost all components of the OptoArm can be substituted for cheaper alternatives that may be readily available. The mounting and optical components (Table 1) can be easily interchanged with alternatives. The required components related to thermal management, however, are highly dependent on the requirements of the electronical components (see *thermal stability* section). Moreover, the choice for the specific LED-driver in this paper is also related to the possible automation of the system, which we will touch upon in later sections. In the end it comes down to the user to verify that any alternative components meet the required specifications of the system and that the system still performs adequately for the desired experimental set-up.

#### Thermal management

Although LEDs rank among the most efficient sources of illumination a large portion of the input power is still converted to heat (Lin et al., 2011; Fang et al., 2017). It is important to have adequate thermal management to remove this waste heat by either conduction or convection, as excessive increases in temperature at the LED junction directly affect LED-performance. In a short-term this could result in color shifts and reduced light output (intensity), while in the long-term accelerated lumen depreciation may take place (Park et al., 2005; Lee et al., 2011; Huang et al., 2015; Han et al., 2015; Dalapati et al., 2016). While the so-called “T-droop”, i.e. the efficiency droop with increasing temperature, is more pronounced in AlGalnP-based LEDs (e.g., red LEDs); blue InGaN-LEDs still suffer from (minimal) color shifts and decreased output due to high junction temperature (Kim, et al., 2016; Fu et al., 2018; Oh et al., 2019). The junction temperature of an LED is determined by three parameters: drive current, heat dissipation and the ambient temperature (Kim et al., 2016). While the drive current can be kept stable with fixed-current drivers and ambient temperature can be controlled for in climate-controlled rooms, efficient heat removal generally requires extra management. Heat is typically transferred away and dissipated from the LED using passive systems like heat sinks (Figure 2F). The required thermal properties of such a heat sink can be calculated by using a basic thermal model: see box 1. We selected a 5.2 °C/W heatsink that fulfills our criteria for a system with appropriate thermal cooling. It is important to underline that the measured junction temperature is higher than theoretically expected when using a heatsink of 5.2 °C/W (see box 1). Therefore, it is critical to use a safe margin when selecting a heatsink and to verify the actual junction temperature instead of simply relying on the thermal equation.

### Lighting area and intensity is stable and can be adjusted to experimental requirements through exchangeable lenses and adjustable working distance

Next, we explored the usability of the OptoArm for experimental purposes with *C. elegans*. We evaluated different aspects of the system that are critical for optogenetics, starting with light intensity and stability. The Royal blue Luxeon Rebel LED of the OptoArm has a beam angle of 125°, which results in a large surface being illuminated. As intensity is the function of power divided by surface, there are simply two parameters that determines the intensity of a system: 1) area being illuminated, 2) the power of the LED. The maximal power of the LED (1.03 W) at a current of 700 mA is fixed and cannot be further enhanced. However, the area that is illuminated can be reduced by either decreasing the working-distance (WD) or applying convex optics to concentrate the light beam to a smaller surface. Since optogenetic experiments in *C. elegans* require an intensity of at least 1.0 mW/mm^2^ and preferably 1.6 mW/mm^2^ (Nagel et al., 2005; Liewald et al., 2008; Schmitt et al., 2012; Husson et al., 2013), we explored the produced intensity of the OptoArm without and with several low-cost lenses (Figure 3A, B).

**Figure 3:**
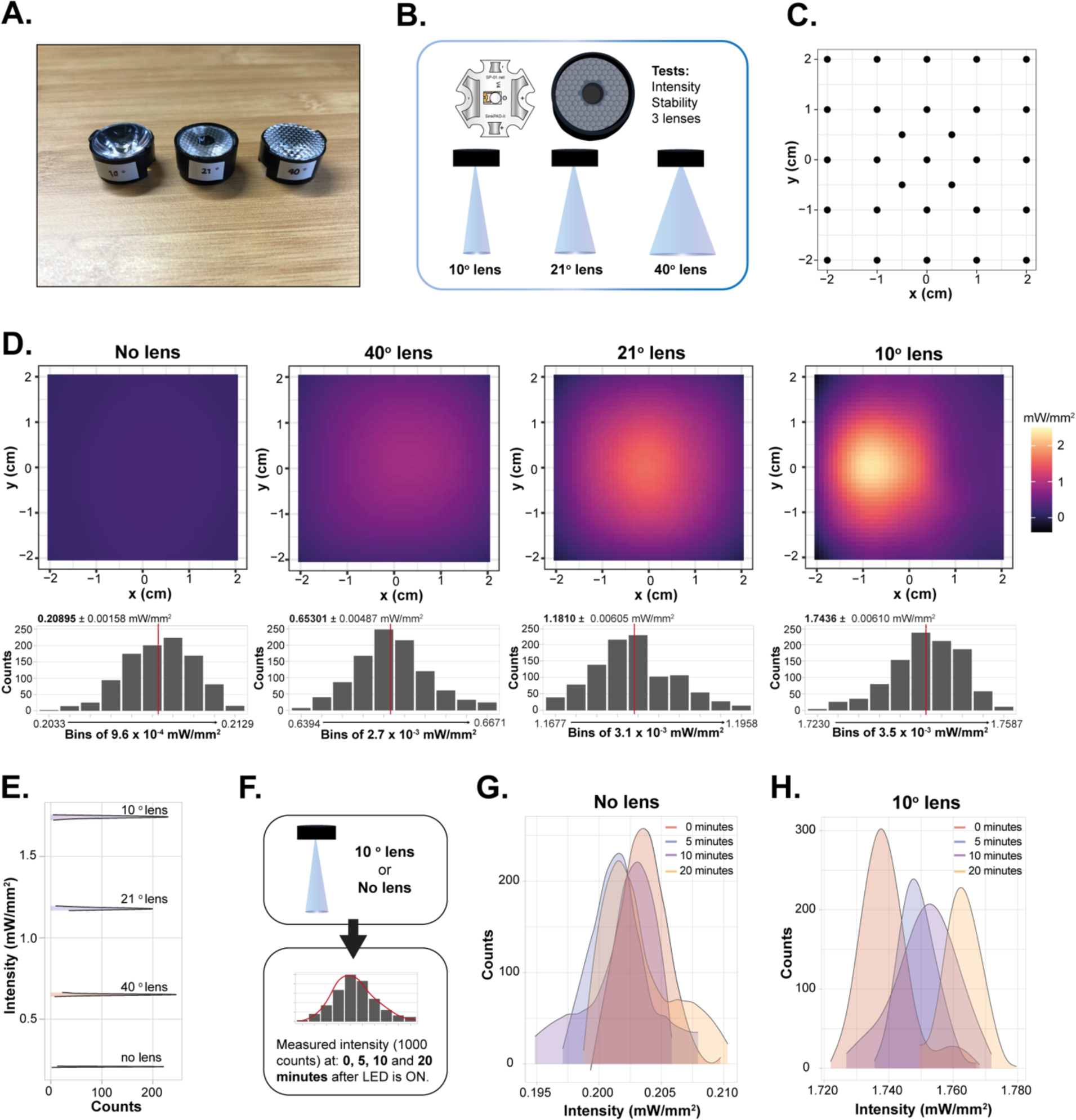
The OptoArm provides the light intensity and stability required for optogenetic experiments. **A)** The different lenses used for testing. **B)** A schematic outlining the tested parameters with the three different lenses. **C)** Raster showing the *x,y* coordinates used to measure the intensity of the LED. **D)** The different integrated intensity profiles of the LED with and without lenses. The histograms show the zoomed-in distribution of intensity-readings between a minimum and maximum intensity with a specified bin width. Red lines show the average intensity. **E)** Smoothened histograms of D that show the increase in intensity when using different lenses with the OptoArm. **F)** Schematic showing the method to assess the stability of intensity over time. **G)** Distributions of intensity readings at the different time-point without a lens **H)** and with a 10° lens. All readings were performed at 448nm.

To compare the different set-ups, we used a fixed working-distance of 5 cm between the LED and the surface to be illuminated. With a power meter set to 448 nm, we measured the light intensity of 29 different *x,y* coordinates in an area of 4 by 4 cm using a ‘hot-spot’ approach (Figure 3C). From those collected measurements we generated integrated intensity profiles (Figure 3D). We observed clear differences in the acquired intensity profiles between the different set-ups. Without a lens, the illumination is relatively uniform, while using lenses that narrow the angle of the light beam result in more pronounced gaussian-like profiles with clear intensity peaks in the middle. The combination of the OptoArm with a 10° lens fulfills the required intensity of 1.6mW/mm^2^. In order to assess the stability of these profiles, we collected 1000 (∼ 1 minute) readings per set-up at the *x,y* coordinate with the highest intensity, thereby assuming an gaussian-distributed light source. For all the set-ups, the readings were steady and consistent, as evidenced by the narrow distributions (Figure 3D, 3E). Clearly, adding different convex lenses to the OptoArm can modify the output of the system in terms of measured light-intensity (Figure 3E). Note that the readings in the histograms (Figure 3D, E) are lower than those seen in the intensity profiles (Figure 3D). The latter can be explained by the different measuring methods (software-related) that were used: hot-spot detection versus a gaussian assumption.

Although most optogenetic experiments only take a few milliseconds, or a minute at its maximum (Busack et al., 2020), we also assessed the stability of the system in terms of intensity over several minutes (Figure 3F). With the same working-distance of 5 cm, we assessed the maximum intensity at 0, 5, 10 and 20 minutes after turning the LED on. Again, we collected 1000 readings per timepoint. The ON-time of the LED had no influence on the measured intensity without a lens (Figure 3G), while only a minor drift in intensity (<0.02 mW/mm^2^ in 20 minutes) was observed with the 10° lens (Figure 3H). Therefore, we conclude that the OptoArm, in its current set-up, performs well in terms of intensity and stable output over time.

### Due to its compact and highly flexible nature, the OptoArm is compatible with both single- and multi-worm imaging platforms

When using the OptoArm with a microscope or any tracking platform (e.g. the WF-NTP), the angle of incidence of the blue light is in most cases not perpendicular to the surface as a consequence of practical limitations, e.g. the arm should not block the light-pad from the sample to the ocular or camera (Figure 4A). The consequence of working under a non-perpendicular angle is that intensity profiles will deform due to increases and decreases of the path length of specific parts of the light beam (Figure 4B, C). The intensity of light as a function of the distance from the light source follows an inverse square relationship. Therefore, increased path lengths will result in a significant reduction of the light intensity.

**Figure 4:**
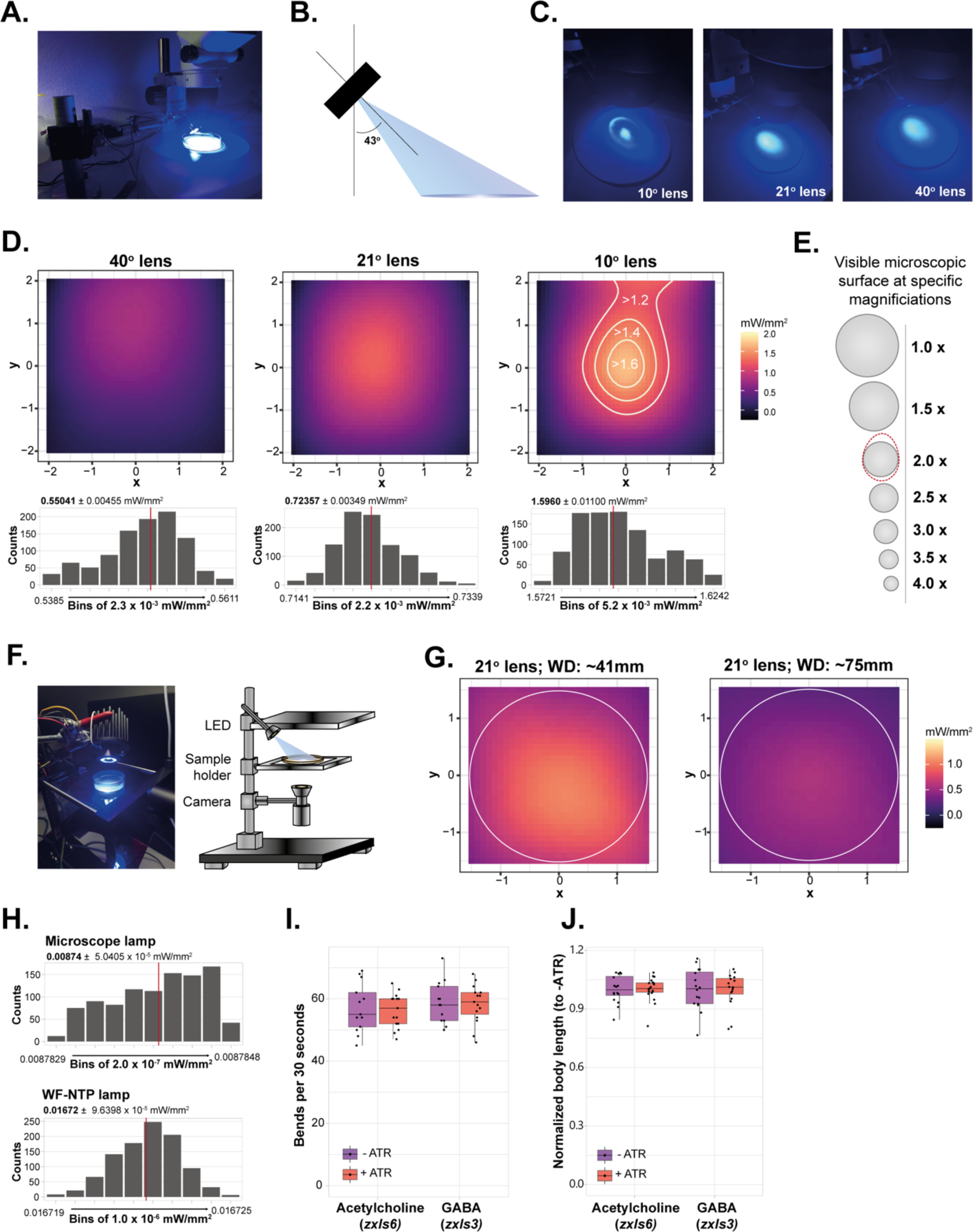
The OptoArm fulfills all technical criteria in different experimental set-ups. **A)** The OptoArm set-up used with a standard dissection microscope. **B)** In experimental set-ups, the OptoArm is used with a specific angle (43 °), thereby deviating from perpendicular illumination. **C)** The different illumination profiles for the different lenses upon and illumination with the specified angle of incidence. **D)** The different integrated intensity profiles for C. The histograms show the distribution of intensity-readings between a minimum and maximum intensity with a specified bin width. Red lines show the average intensity. **E)** Schematic showing the different visible microscopic areas (of the used microscope) when using different magnifications. The areas are on scale and can be directly compared to the intensity maps shown in D. 1.0x corresponds to a total magnification of 15x, and 4.0 to a total of 60x. The dotted red line represents the area in D with an intensity of >1.6 mW/mm^2^. **F)** The OptoArm set-up with the WF-NTP. **G)** The different integrated intensity profiles of the LED with the 21° lens, at different working distances with a fixed angle of incidence of 43°. Profiles were generated in the set-up with the WF-NTP. Circles represent the outline of a 3-cm NGM plate. **H)** Histograms of intensity-readings of the microscope-light (gaussian) and WF-NTP back-light (flat-field). Red lines show the average intensity. All readings were performed at 448nm. **I)** Thrashing frequency of D1 worms, expressing ChR2 under the *unc-17* (acetylcholine) or *unc-47* (GABA) promotor, recorded with the WF-NTP, grown with or without ATR. There is no significant effect of the WF-NTP light on this behavior, *n = 15*, two-tailed unpaired Student’s t test (all: n.s.). **J)** Body length of D1 worms recorded with the WF-NTP, grown with or without ATR. Body length was normalized by the mean the of the paired ATR-condition. There is no significant effect of the WF-NTP light on this behavior, *n = 15*, two-tailed unpaired Student’s t test (all: n.s.). For **I)** and **J**) there was no photostimulation with the OptoArm.

To validate the OptoArm in an actual experimental set-up, we again generated integrated intensity profiles and stability measurements of the three lenses when used with a standard dissection microscope (Figure 4). As expected, both the hot-spot intensity and the steady-state measurements pointed out that the maximal intensity is decreased when working under a non-perpendicular incidence angle (Figure 4D). However, the acquired intensity with a 10° lens still reaches the required 1.6mW/mm^2^ (WD: ∼3.5 cm) and with some fine-adjustments in height (i.e. working distance) and angle, the intensity even exceeds this number (see also Figure 5). More importantly, the area with the required intensity (1.6mW/mm^2^) is large enough to ensure the whole visible microscopic surface being illuminated with the same intensity (see Figure 4E for details). In fact, an area with a ∼*φ* 1 cm appears to fulfil the required 1.6 mW/mm2 (Figure 4D) and this area corresponds with the visible microscopic surface at a total magnification of 30 x (Figure 4E, 2.0x).

**Figure 5:**
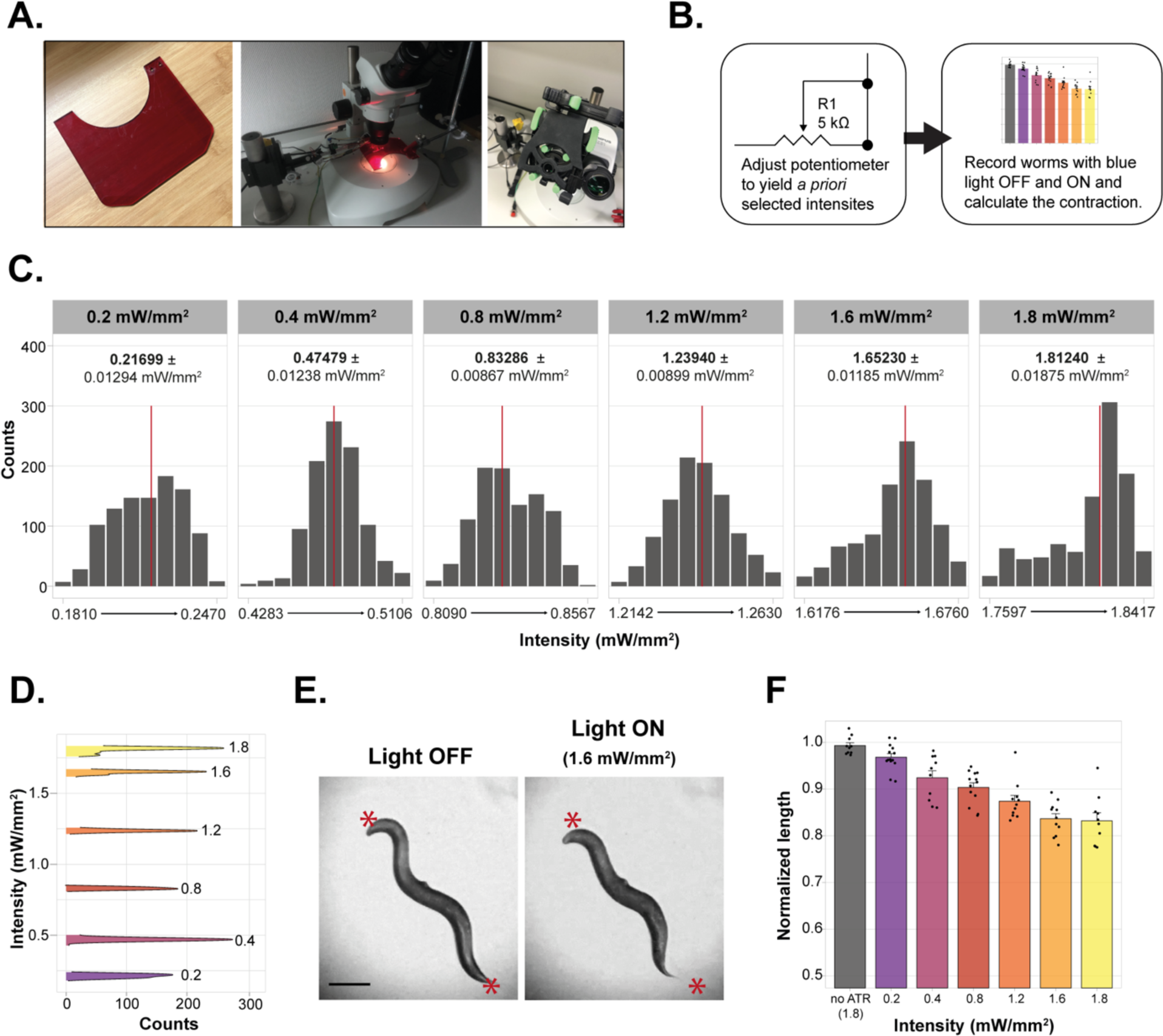
The OptoArm allows manual intensity-adjustments, which modulate biological readouts. **A)** To record worms in an inexpensive way, we used a Carson smartphone adapter to place and orient a cellphone camera to a microscope. A blue-light filter was placed between the ocular and sample to avoid interference of the blue light with recording. **B)** Schematic of the experimental outline: *a priori* determined intensities (0.2, 0.4, 0.8., 1.2., 1.6 and 1.8 mW/mm^2^) were set by adjusting the resistance of the potentiometer. At the different intensities the Δ body length was measured**. C)** The histograms show the distribution of intensity-readings at *a priori* specified intensities. Red lines show the average intensity. **D)** Smoothened histograms of C that show the increase in intensity when adjusting the resistance of the potentiometer. **E)** Pictures of a D1 worms (*myo-3p::ChR2::YFP)* just before and during illumination with 1.6 mW/mm^2^ blue light. Scale bar: 200 *u*m. **F)** The different body lengths (normalized to length before illumination) at different intensities of illumination with the OptoArm. Calculated via a midline-approach, *n =* 10.

Having established and validated a microscope set-up, we moved on to an experimental set-up involving a multi-worm tracker (Figure 4F). When working with a platform that allows multiple worms to be tracked at the same time, it is critical that illumination and light intensity are as uniform as possible at all *x,y* coordinates in the region of interest (ROI). Spatial consistency is relevant in order to reduce individual variation. Clearly, lenses that generate narrower gaussian-like profiles (Figure 4D) do not meet those criteria, even though they do provide the adequate intensity. Consequently, we only generated intensity profiles of the OptoArm with the 21° lens at two different working-distances (Figure 4G) and a fixed incidence-angle of 43°. While far from perfect, the 21° lens is offers the best compromise between uniform illumination and high intensity (0.75-1.0 mW/mm^2^) when using 3-cm worm plates (Figure 4G). Moreover, systems like the WF-NTP make it possible to only analyze worms in a specific ROI, which would make it possible to account for spatial inconsistency of illumination at the edges of a plate. If the experimental conditions allow it, one can also simply use smaller plates, e.g., 12-wells, to better fit the intensity profiles. Finally, it should be underlined that the adaptability of the OptoArm makes it easy to further enhance the system e.g. by using triple-LED systems instead of single LEDs to triple the power of the system (Table 1) if required.

Finally, we assessed the intensity of backlights of a standard microscope (maximal intensity) and the WF-NTP (Figure 4H). While, the intensity of those lights is relatively low, it might be sufficient to induce optogenetic responses (Schultheis et al, 2011a). We have tested this hypothesis by comparing the thrashing capacity (i.e. the frequency of lateral bends in liquid) and body length of worms grown with or without ATR with the WF-NTP. Since ATR is required for functional ChR2, non-ATR worms will not respond to blue light and can be used as negative controls (Nagel et al., 2005). We did not find clear evidence for a light-induced response when performing optogenetic experiments with the WF-NTP backlight. Worms grown with or without ATR had a similar thrashing capacity and body length (Figure 4I, J). However, we would recommend to substitute the WF-NTP backlight with a red-shifted variant of 625 nm (Edmund Optics, Red advanced illumination side-fired backlight, #88-411) or to filter out the blue light specifically. This is advisable because several neuronal mutants have been shown to have an increased sensitivity to low-light conditions (Liewald et al., 2008). For microscopic set-ups we suggest to turn down the backlight as much as possible, without compromising the contrast between the worm and its background (Figure 4H).

### Adjustable intensity levels enable titration of the biological response

All *C. elegans* neurons that have previously been analyzed by photo-electrophysiology exhibited graded transmission. Herein, the release of neurotransmitters directly correlates with the extent of depolarization evoked by light-induced ChR2 activation (Husson et al., 2013). This type of transmission has clear consequences for optogenetic experiments, as changing the light intensity will likely also change transmitter release. Therefore, to properly compare different interventions or different mutants with each other, light intensity should always be consistent and stable throughout complete experiments. On the other hand, the graded transmission in *C. elegans* also allows the ‘titration’ of the extent of ChR2-induced depolarization by adjustment of the light intensity. In this way, one can modify transmitter release and subsequently also fine-tune the behavioral output (Liewald et al., 2008; Husson et al., 2012a).

We explored the adjustability of the OptoArm for ‘titration’ purposes (Figure 5). First, we created a basic microscopic set-up in which a microscope-based smartphone adapter was used to allow movies being made with a smartphone instead of an expensive camera (Figure 5A). This set-up was completed with a standard blue-light/UV-protection shield to avoid saturation of the camera. Secondly, we used the potentiometer of the OptoArm to adjust the intensity of the system to specific values, without changing the WD or angle of incidence (e.g., 0.2; 0.4; 0.8; 1.2; 1.6 and 1.8 mW/mm^2^) (Figure 5B). Then, we measured both the stability of each specified light intensity and the accompanying body length of worms expressing ChR2 under the *myo-3* promoter before and during illumination (Figure 5C-F). The normalized body length was then calculated by dividing the body length during illumination by the body length before illumination. We were able to adjust the potentiometer in such a way that the *a priori* specified intensities could be acquired relatively well and were consistently stable over 1000 readings (Figure 5C, D). Clearly, adjusting the resistance of the potentiometer of the OptoArm can modify the output of the system in terms of measured light-intensity (Figure 5D). The different light-intensities had dose-dependent effects on normalized body length up till 1.6 mW/mm^2^ (Figure 5E,F). Taken all together, the OptoArm allows users to change the light intensity and thus to study its effect on behavioral changes.

### Perimeter and midline measurements are highly correlated and can be used interchangeably as a readout for body length

Before continuing with the biological validation of the OptoArm, we first took the time to explore the possible methods to quantify changes in body length during an optogenetic experiment. In general, to follow changes in body-length over time, one can manually draw a midline through the worm from head to tail before and during blue-light stimulation (Figure 5). Clearly, this labor-intensive scoring method is vulnerable to inconsistency and experimenter bias and becomes impractical when scoring many worms and conditions. There are Matlab codes available to automate and streamline these measurements (Liewald et al., 2008; Crawford et al., 2020), but these codes also require a basic understanding of programming languages and access to Matlab. Based on the idea that the OptoArm should be a simple system that is accessible to the whole community, we aimed to simplify the accompanying analysis for ‘Δ body length’, i.e. the change in body length upon optogenetic stimulation, as well. Therefore, we explored the use of the opensource software ImageJ to analyze the optogenetic data in a semi-automatic way (Figure 6).

**Figure 6:**
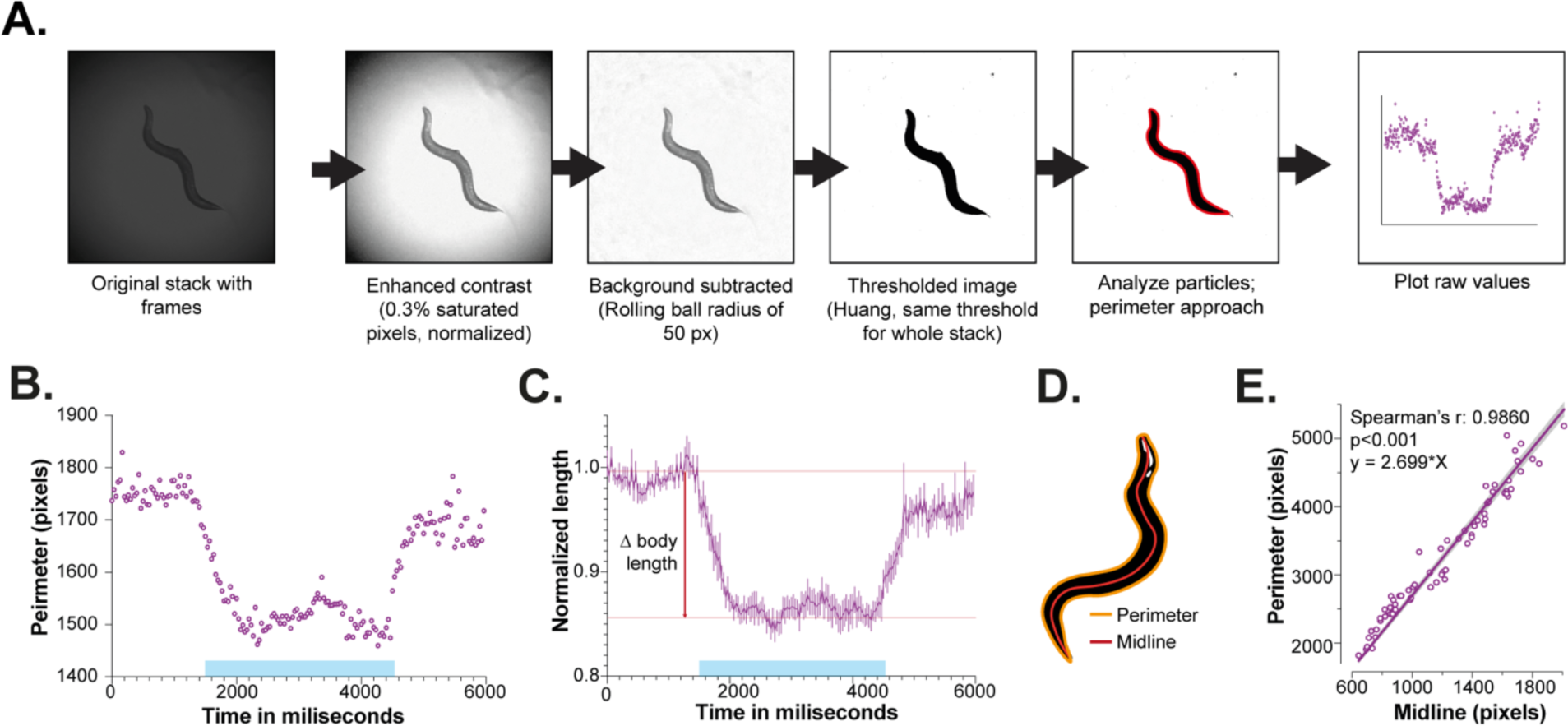
A semi-automatic perimeter-approach is a reliable way of assessing changes in body length during and after optogenetic stimulation. **A)** Schematic outline of image-processing in ImageJ to follow body-length changes over time in a semi-automatic way. **B)** Raw readings of body length (perimeter in pixels) of a worm expressing ChR2 in cholinergic neurons (*unc-17p::ChR2::YFP)* plotted against time in milliseconds, the graph represents n = 1. **C)** The change in body lengths (normalized to length before illumination) of multiple worms, *n = 10*, when using a perimeter approach to estimate the worm length. Δ body length equals the change in length before illumination and during illumination. **D)** Schematic of a worm with the midline and perimeter highlighted. **E)** The relationship between the midline of the worm and the perimeter. *n = 70* (35 still images of worms at light ON and 35 worms at light OFF). The spearman correlation was calculated (p<0.001).

It has been stated that the perimeter of the worm is equal to two times the body length plus two times the diameter (Kammenga et al., 2007). Since the perimeter can be easily measured and is more often used as a way to represent body length in *C. elegans* (Fujiwara et al, 2002; Kammenga et al., 2007; Nussbaum-Krammer et al., 2015), we created a sequential pipeline in ImageJ with the perimeter as readout (Figure 6A). All steps of this ImageJ pipeline, and the accompanied methods, are included in the material and method section. In short, the pipeline focusses on enhancing contrast, subtracting background and thresholding to yield a set of binary images from the worm without gaps or ruffled edges. The binary images are used to measure the perimeter of the worm in each frame. Since all steps are performed on complete movies, the end-result is a list of perimeters for a single worm over time. In Figure 6B, the raw perimeter measurements of a worm expressing ChR2 under the *myo-3* promoter, before, during and after optogenetic stimulation are plotted. By normalizing all individual measurements of a single worm to the average perimeter in the first 60 frames (light OFF) one can create kinetic-plots of the change in perimeter that represent multiple worms (Figure 6C). In order to validate this method, we measured the length of the midline and the accompanying perimeter of 70 individual worms (Figure 6D). The midline and perimeter correlate significantly and therefore appear to be both good estimates of body length (Figure 6E).

### Optogenetic stimulation of cholinergic and GABAergic mutants with the OptoArm accurately reproduces the biological response observed with a more expensive system

To validate the OptoArm and accompanying image-processing biologically, we aimed to reproduce previously described and published optogenetic data on neuronal mutants (Figure 7A). It has been showed that mutations in the *unc-47* gene, which represents the vesicular GABA transporter vGAT (McIntire et al., 1997), translate into the lack of body wall relaxation upon optogenetic stimulation of GABAergic neurons, but increased contractions when cholinergic neurons were photo-stimulated (Liewald et al., 2008). At the same time, mutations affecting the cholinergic system, e.g. *unc-26* that represents the phosholipid phosphatase synaptojanin (Figure 7A), often translate into significantly stronger body-wall muscle contractions upon cholinergic stimulation as well. The latter was initially marked as contradictory as electrophysiological data of those mutants showed reduced electrically evoked postsynaptic currents (ePSCs). However, the paradoxical results are most likely a result of compensatory mechanisms in the muscle, underlining the complementary information that behavioral optogenetic experiments can provide (Liewald et al., 2008).

**Figure 7:**
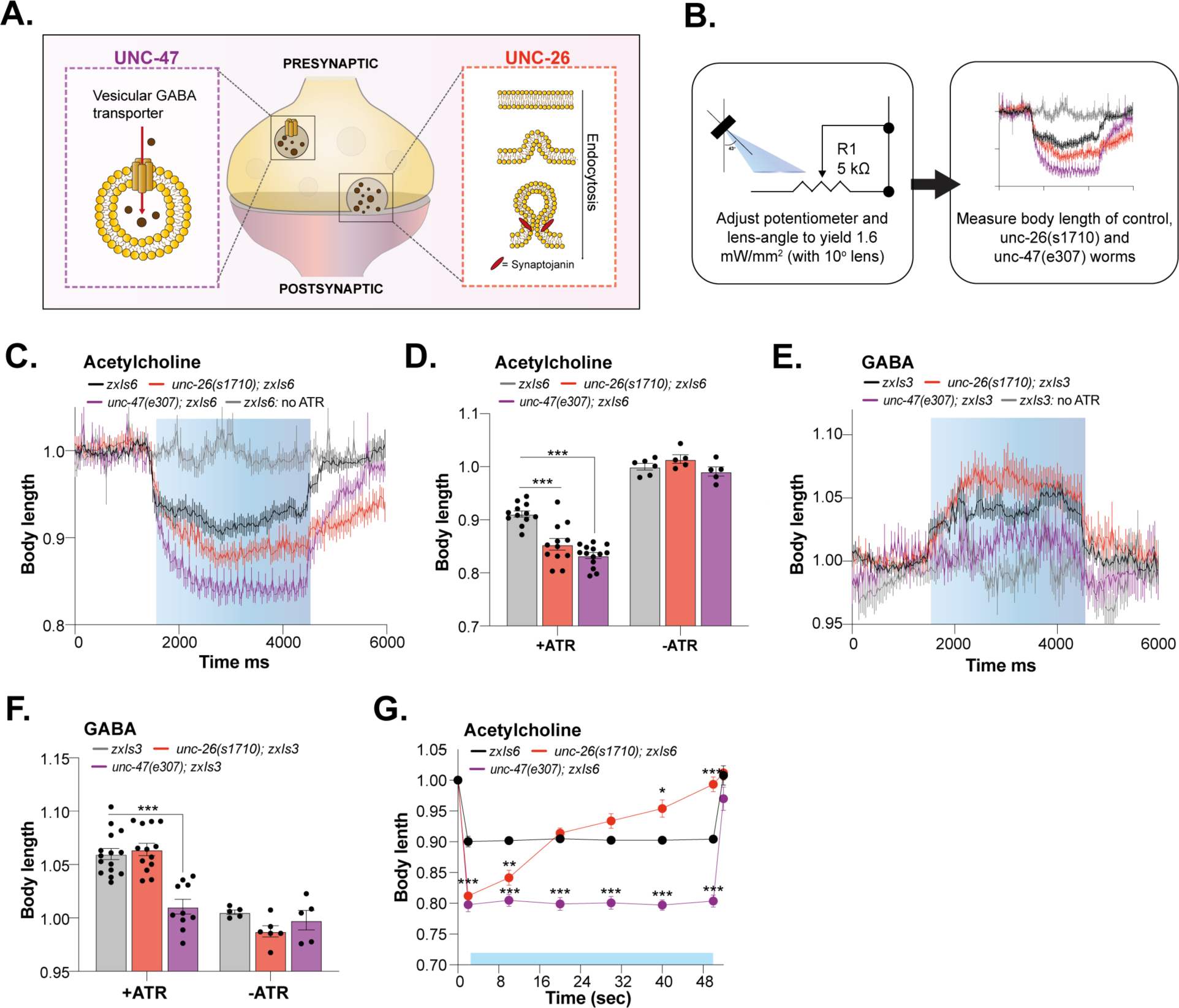
Mutations in the cholinergic or GABAergic system affect **Δ** body length after optogenetic stimulation with the OptoArm. **A)** A schematic showing the different mutant used to verify previously established results. **B)** Schematic of the experimental outline: the change in body length was measured after blue light stimulation with an intensity of 1.6 mW/mm^2^. **C)** Kinetics of optogenetic stimulation in mean ±SEM body length of control worms and *unc-26(s1710*) and *unc-47(e307)* mutants expressing ChR2 in cholinergic neurons, *n = 9-10*. A perimeter-approach was used. **D)** Quantification of mean ±SEM normalized body length of control D1 worms and *unc-26(s1710)* and *unc-47(e307)* mutants expressing ChR2 in cholinergic neurons. Two-way ANOVA (Interaction, ATR, Genotype: p<0.001) with post-hoc Dunnett, *n = 11-15* for ATR+ and *n = 5-6* for ATR-. A midline-approach was used. **E)** Kinetics of optogenetic stimulation in mean ±SEM normalized body length of control worms and *unc-26(s1710)* and *unc-47(e307)* mutants expressing ChR2 in GABAergic neurons, *n = 9-11.* A perimeter-approach was used. **F)** Quantification of mean ±SEM normalized body length of control worms and *unc-26(s1710)* and *unc-47(e307)* mutants expressing ChR2 in GABAergic neurons. Two-way ANOVA (Interaction, ATR, Genotype: p<0.001) with post-hoc Dunnett, *n = 10-15* for ATR+ and *n = 5-6* for ATR-. A midline-approach was used**. G)** Long-term photostimulation of control worms, unc-26(s1710) and unc-47(e307) mutants expressing ChR2 in cholinergic neurons. Two-way repeated ANOVA with Geisser-Greenhouse correction (time, genotype, time x genotype and individual worms: p<0.001) and post-hoc Dunnett, *n = 10*. A midline-approach was used. Blue bars represent ‘light ON’. Acetylcholine: *zxIs6 (Punc-17::ChR2::YFP), zxIs3 (Punc-47::ChR2::YFP).* All experiments were replicated three times, one representative experiment is shown. *: p ≤ 0.05, **: p ≤ 0.01, ***: p ≤ 0.001. Error bars represent S.E.M.

Here, we explored two mutants that have been described in the specified paper, namely *unc-47(e307*) and *unc-26(s1710)*. We used a microscopic set-up (Figure 5A) combined with the OptoArm, thereby ensuring a light intensity of 1.6 mW/mm^2^ (Figure 7B). Next, D1 adults expressing ChR2 specifically under the *unc-17* promoter (i.e. expression in cholinergic neurons; *zxIs6*), grown on ATR- or ATR+ plates, were exposed to a regime of 3 seconds light off – 3 seconds light on – 3 seconds light off while being recorded. Subsequently, the movies were used to determine the body length over time by a perimeter-approach (Figure 7C) or by measuring the length at fixed timepoints before (500 ms before light ON) and during illumination (1000 ms after light turned ON) with a midline-approach (Figure 7D). Similar to the results described by Liewald et al., we found that *unc-47(e307)* and *unc-26(s1710)* mutants showed significantly stronger photo-evoked contractions (Figure 7C), with clear differences in muscle-relaxation kinetics after the photo-stimulation was terminated. In addition, worms raised at -ATR plates, did not respond to blue light with body contractions and served as appropriate negative controls. The change in body length of the *unc-47* (mean ± *SD*: 0.83 ± 0.020), and *unc-26* mutants (0.85 ± 0.037) and control (0.91 ± 0.02) worms, as a result of photo-stimulation with the OptoArm, are similar to the numbers found by Liewald et al (2008) (WT: ∼0.92, *unc-26*: ∼0.87, *unc-47*: ∼0.86) (Figure 7D). Next, we also explored the effects of GABAergic stimulation in the *unc-47(e307*) and *unc-26(s1710)* mutants, by expressing ChR2 under the *unc-47* promotor (*zxIs3*). The same lightening regime was used and the body length was again determined by a perimeter-approach (Figure 7E) and a midline-approach (Figure 7F). As expected, *unc-47(e307)* mutants (1.01 ± 0.022) lacked a clear relaxation response, while *unc-26(s1710)* worms (1.06 ± 0.020) behaved much more similar to control worms (1.06 ± 0.020), with the exception of a higher initial peak of relaxation (Figure 7E, F). The same observations were described by Liewald et al., (2008) (WT: ∼1.04, *unc-26*: ∼1.00, *unc-47*: ∼1.04).

It has been shown that characteristic defects in neurotransmitter recycling can be reflected by aberrations in long-term photo-stimulation experiments with worms expressing ChR2 under the *unc-17* promoter. Synaptojanin (*unc-26*) is required for endocytotic recycling of synaptic vesicles and mutants are defective in synaptic vesicle budding and uncoating during clathrin-mediated endocytosis (Figure 7A) (Harris et al., 2000; Schuske et al., 2003). The inability to recycle vesicles results in early fatigue of neurons that could potentially translate in the inability to sustain contraction of body wall muscles. Indeed, Liewald et al. (2008) described that wild-type worms normally sustain constant contraction throughout long-term illumination (>60 seconds), while the initially exaggerated body contraction of *unc-26(s1710)* mutants returns quickly to unstimulated levels. We validated this biological data with the OptoArm by following individual worms for 52 seconds, in which worms were constantly illuminated from second 1 to 51 (Figure 7G). While control worms and *unc-47(e307)* mutants showed sustained contraction over the complete interval, *unc-26(s1710)* mutants lost their contractibility already after 8 seconds and returned to wild-type levels after 15 seconds. At the end of the illumination period, the *unc-26(s1710)* worms returned to their initial non-illuminated length and thus lost their contraction completely (Figure 7G). Altogether, we validated the OptoArm by showing that the inexpensive system can reproduce biological findings acquired with a more expensive fluorescence microscope. The use of the OptoArm with ImageJ provides a sensitive platform to study the various functional aspects related to body contraction, elongation and the kinetics of this behavior.

### Combination of the OptoArm with various set-ups allows the readout of multiple biological parameters for a more comprehensive analysis

Having a technically and biologically validated system, we continued exploring the use of the OptoArm for different applications and set-ups. Since the platform also allows multi-worm illumination (Figure 4), we explored the use of the OptoArm together with a multi-worm tracking platform, in this case the WF-NTP (Perni et al., 2018; Koopman et al., 2020). It has been previously shown that photo-stimulation of worms on solid substrates, expressing ChR2 under the *unc-47* promotor (GABAergic neurons) or under the *unc-17 promotor*, reduces locomotion speed (Liewald et al., 2008; Kittelmann et al., 2013). The observed decline in locomotion speed is caused by different mechanisms. The photoactivation of ChR2 in GABAergic neurons induces a flaccid paralysis, during which worms straighten up completely for a few seconds (Liewald et al, 2008). This observation is consistent with the inhibitory role of GABA in the neuromuscular system. Stimulation of cholinergic neurons, on the other hand, causes a spastic paralysis during which worms display dorsal coiling behavior in addition to body-wall muscle contraction (Liewald et al., 2008; Kittelmann et al., 2013). This coiling behavior can be quantified by following changes in eccentricity. Eccentricity is used as a measure of how nearly circular an ellipse is. Typically, crawling and thrashing worms have an average eccentricity close to 0.93 or higher. At the same time, extreme coilers can have an average eccentricity close to 0.6-0.7 (Koopman et al., 2020).

In liquid, photo-stimulation of ChR2 in GABAergic neurons inhibits swimming behavior, quantified as the decline in trashing frequency (Liewald et al., 2008). The exact effects of cholinergic stimulation on swimming behavior, however, still need to be determined. Both swimming behavior and coiling behavior in liquid can be assessed with the WF-NTP (Koopman et al., 2020), allowing other readouts, in addition to changes in body length, to be collected. Therefore, we analyzed thrashing frequency and eccentricity of worms in liquid expressing ChR2 under the *unc-17* or *unc-47* promotor to evaluate population-based optogenetic experiments with the OptoArm and the WF-NTP (Figure 8A). We first explored the thrashing ability of worms grown with and without ATR before and during photo-stimulation. Photoactivation of ChR2 causes a decrease in thrashing behavior independent of which neuron-population was stimulated (Figure 8B). The kinetics of the observed movement-decline was however strikingly different (Figure 8C). GABAergic stimulation causes an initial decrease in thrashing behavior that gradually fades back to normal levels, while the cholinergic effect appears to saturate at the end illumination-period (Figure 8C). Indeed, it has been shown before that worms recover partially from paralysis even tough illumination of GABAergic ChR2 is sustained (Liewald et al., 2008). Taking this difference in kinetics into account, we only analyzed the first 10 seconds of the movies for the worms with GABAergic ChR2-expression and the last 10 seconds for worms with cholinergic ChR2-expression (Figure 8D). While the decline in thrashing behavior was already evident and significant when complete movies were analyzed (Figure 8B), the effect was more pronounced when taking the underlying kinetics into account (Figure 8D). Clearly, the population effect for GABAergic stimulation is most prominent directly after photo-stimulation while the cholinergic system requires more time for saturation of the effect. Importantly, at a blue-light intensity of 1 mW/mm^2^, photoactivation of ChR2 in cholinergic-neurons does not result in complete paralysis of worms in liquid (Figure 8B-D). However, we did observe clear decreases in eccentricity when cholinergic neurons were stimulated via ChR2-based illumination (Figure 8E), while the eccentricity was unchanged after GABAergic stimulation or when worms were grown without ATR (Figure 8E,F). Again, we did observe that this effect could be maximized by long-term stimulation and analysing the last 10-seconds of the assay. Interestingly, coiling behavior upon cholinergic stimulation has been ascribed to the concomitant GABA release that is indirectly triggered by photoactivation of cholinergic neurons innervating GABAergic neurons (White et al., 1986; McIntire et al., 1993; Schuske et al., 2004; Liewald et al., 2008). Coiling behavior is completely absent when GABAergic mutants expressing ChR2 in cholinergic neurons are illuminated (Liewald et al., 2008). Therefore, coiling behavior likely provides a readout of the interplay between GABAergic and cholinergic neurons within the nervous system.

**Figure 8:**
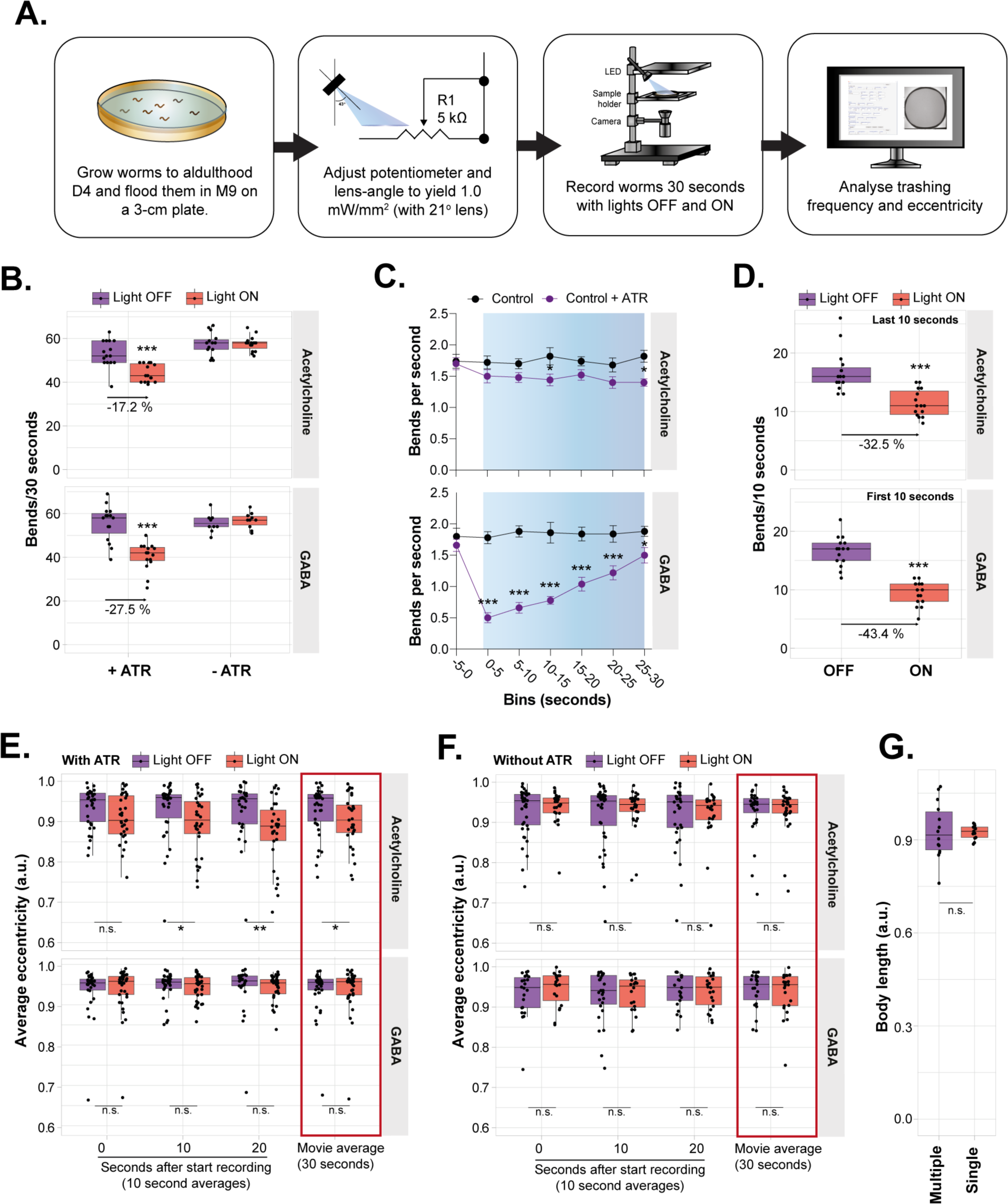
Population characteristics acquired with the WF-NTP are potential readouts of optogenetic stimulation with the OptoArm. **A)** Schematic of experimental outline. D4 worms were assessed in liquid with the OptoArm being OFF and ON and population characteristics (bending frequency and eccentricity) were analyzed with the WF-NTP software. **B)** Change in bends per 30 seconds when light is ON of worms grown with or without ATR. The percentual decline in thrashing capacity is annotated in the graph, *n = 15.* For acetylcholine: Mann Whitney U test p<0.001, GABA: two-tailed unpaired Student’s t test: p<0.001. **C)** Binned effect of blue-light on swimming behavior of transgenic worms grown with or without ATR, *n = 10*. Two-way ANOVA (time, genotype, interaction: p<0.001) with post-hoc Sidak’s. **D)** Change in bends per 10 seconds after optogenetic stimulation. Acetylcholine: only the last 10 seconds of 30s-illumination is used, GABA: only the first 10 seconds are used. *n = 15.* For acetylcholine: Mann Whitney U test p<0.001, GABA: two-tailed unpaired Student’s t test: p<0.001. **E)** Effect of blue-light on eccentricity of worms grown with or **F)** without ATR. *n = 20 – 40,* Mann-Whitney U tests Acetylcholine + ATR: 0-10s: not significant, 10-20s: p=0.0192, 20-30s: p =0.0033, for 0-30s: p = 0.0427. Acetylcholine – ATR, and both GABA conditions: n.s. **G)** The body length of worms expressing ChR2 under the *unc-17* promoter after blue-light stimulation, *n = 15,* two-tailed unpaired Student’s t test: n.s. Blue bars represent ‘light ON’. Acetylcholine: *zxIs6 (Punc-17::ChR2::YFP), zxIs3 (Punc-47::ChR2::YFP).* All experiments were replicated three times, one representative experiment is shown. *: p ≤ 0.05, **: p ≤ 0.01, ***: p ≤ 0.001.

Finally, we investigated the use of multi-worm illumination and low-resolution (e.g. perimeters of 60-80 pixels instead of 2000-3000 pixels) videos to calculate the body length of illuminated worms expressing ChR2 cholinergically. While the variation between individual worms was much larger when multiple worms were illuminated at the same time, the average length of photoactivated worms was (0.92 ± 0.08) strikingly similar to single-worm illumination (0.92 ± 0.02) (Figure 8G). The variation is most likely caused by the lack of flat-field illumination of the OptoArm and thus, while being small, spatial differences in light intensity (Figure 3 and 4). Taken together, combining the OptoArm with a multi-worm tracker provides a way to perform population-based optogenetic experiments with multiple readouts in addition to body length. The coiling behavior and thrashing capacity are examples of such readouts and provide distinct information about the functionality of the neuromuscular system. However, it should be underlined that many other behaviors and associated readouts can be assessed when ChR2 is expressed in different neuronal populations, e.g., crawling speed for food-slowing (Sawin et al., 2000; possible with the WF-NTP), reversals for touch-neuron stimulation, egg-laying behavior after HSN-stimulation (Leifer et al., 2011) and so on.

### A comprehensive optogenetic analysis reveals a faster age-dependent deterioration in the cholinergic system compared to other components of a neuromuscular unit

The ability to explore neuronal function with optogenetic tools provides an elegant way to study synaptic connectivity and function. As animals age, they exhibit a gradual loss in motor activity (Hosono et al., 1980; Bolanowski et al., 1980; Johnson et al., 1987; Herndon et al., 2002; Huang et al., 2004). Over the years, the hunt for the mechanisms underlying this age-associated decline in movement have been subject to many studies (Hosono et al., 1980; Bolanowski et al., 1980; Johnson et al., 1987; Herndon et al., 2002; Mulcahy et al., 2003; Huang et al., 2004; Liu et al., 2013). For a long time, it has been hypothesized that the age-dependent decline in motor activity is the result of muscle frailty rather than a functional deficit in the nervous system (Herndon et al., 2002). This hypothesis was strengthened by studies showing that the structural integrity of the nervous system was found to be well preserved over time (Dillin et al., 2002; Herndon et al., 2002; Huang et al., 2004; Murakami et al., 2005; Hsu et al., 2009; Kauffman et al., 2010). When other studies described mild morphological deteriorations at synapses at a later age, it was argued that the nervous system also undergoes are-related changed at the morphological level (Pan et al., 2011; Tank et al., 2011; Toth et al., 2012). In 2013, Liu et al questioned whether morphological analysis would be a reliable predictor for the functional status of a tissue. They showed that despite the preservation of integrity, motor neurons undergo an age-related decline in function already starts in early life. At the same time, such a functional decline was absent in body-wall muscles. Thus, they showed that neuronal dysfunction precedes muscle dysfunction (Liu et al., 2013).

While Liu et al., (2013) shed light on the dynamics between the neuronal and muscle system during ageing, they did not specifically focus on the different types of motor neurons. As mentioned earlier, movement in *C. elegans* is the outcome of an antagonistic interplay between GABAergic inhibition and cholinergic stimulation (McIntire et al., 1993; White et al., 1986). The lack of GABAergic input has severe consequence for the movement capacity (Jiang et al., 2005; Tien et al., 2020), while increased GABAergic stimulation has also been shown to results in cessation of thrashing behavior (Liewald et al., 2008; Figure 8). Clearly, the delicate balance between excitation and inhibition determines the capacity to move. While electrophysiology is technically challenging, optogenetics provide a relatively easy way of follow specific neuron populations functionally over time and relate those readouts with behavioral changes. Here, we investigated the age-related decline of both GABAergic neurons and cholinergic neurons specifically and correlated this decline with the deterioration of movement (Figure 9A).

**Figure 9:**
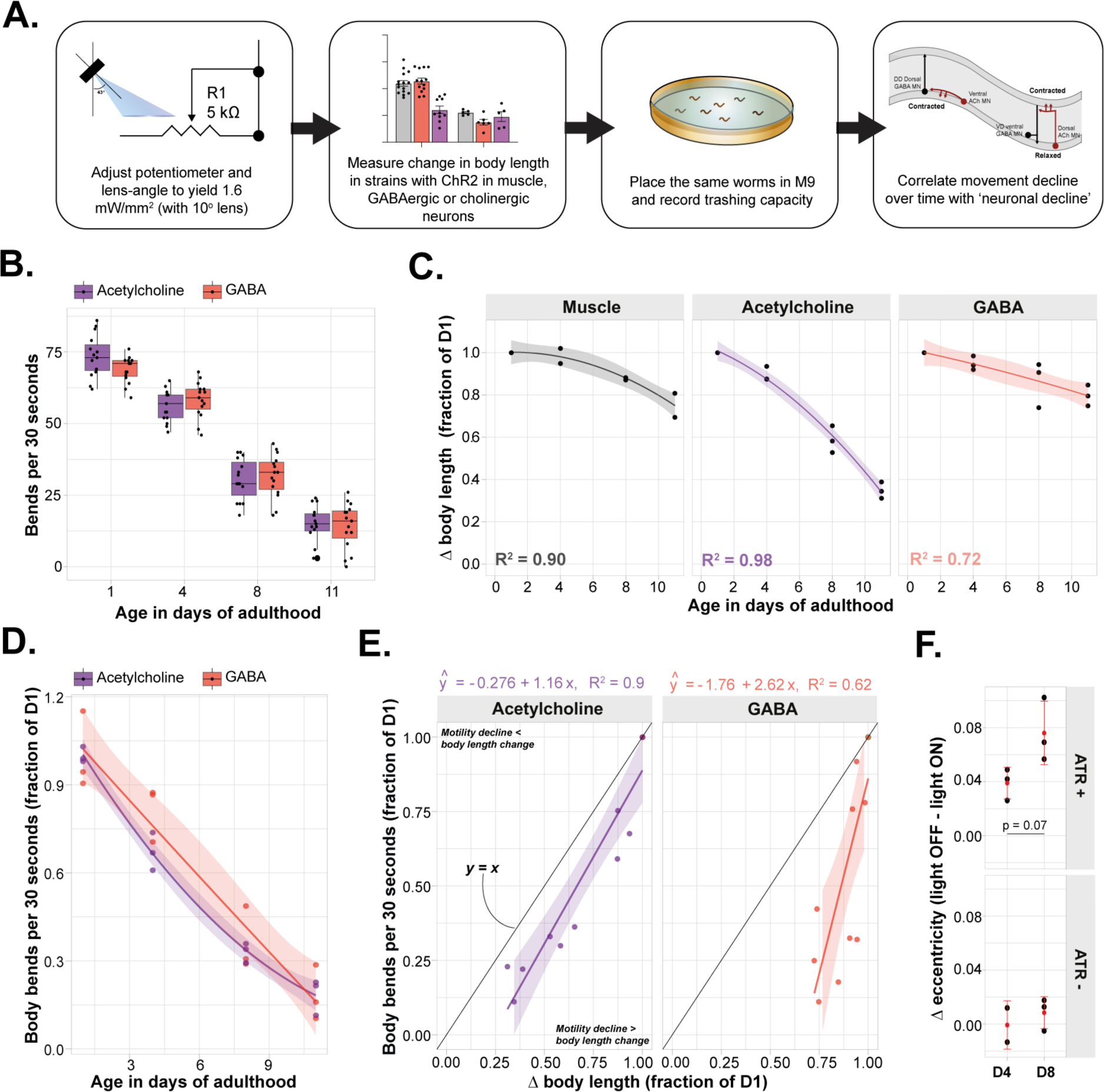
Cholinergic neuronal ageing correlates best with the decline in motility. **A)** Schematic showing the experimental outline. The Δ body length was determined for strains expressing ChR2 in muscle, cholinergic neurons and GABAergic neurons at several time-intervals. At the same time the movement capacity, represented as bends per 30 seconds in liquid, was also measured. **B)** The decline in thrashing capacity in liquid of worms expressing ChR2 in cholinergic of GABAergic neurons. *n = 15*, two-way ANOVA (Age: p<0.001, interaction and neuron type: n.s.). The experiment was triplicated, one representative experiment is shown. **C)** The normalized Δ body length (elongation for GABA, contraction for muscle and cholinergic neurons) at different ages as measured by the midline-approach. Every point represents the mean of an experiment with > 10 biological replicates each. Fitted lines represents a general linear model with 2^nd^ order polynomial fit. All points within one experiment were normalized by the intra-experimental average at D1 within each strain. **D)** The average decline in thrashing ability over time. Every point represents the mean of an experiment with > 20 biological replicates each. All points were normalized by the inter-experimental average at D1 within each strain. This sets the average bends per 30 seconds to exactly 1.0 at D1. Fitted lines represents a general linear model with 2^nd^ order polynomial fit. **E)** The correlation between the Δ body length and Δ movement capacity (normalized to the changes and capacity at D1 intra-experimentally). Fitted lines represent simple linear regression without restrictions (both slopes deviate significantly from zero: p<0.001). The black line corresponds to a relation of *x = y,* in which the decline in movement is equal to the decline in Δ body length. **F)** The Δ eccentricity (eccentricity before illumination minus eccentricity during illumination) at different ages after optogenetic stimulation. Each individual point represents the mean of *n > 15* worms. Student’s t test: n.s. All experiments were replicated three times. The transparent zones around the fitted lines represents the confidence interval (95%). *: p ≤ 0.05, **: p ≤ 0.01, ***: p ≤ 0.001.

We started with assessing the movement decline over time by looking at the thrashing frequency of worms expressing ChR2 under the *unc-17* or *unc-47* promotor. As expected, we observed a significant age-dependent decline in thrashing frequency that is independent of the genotype (Figure 9B). In order to study the relation between neuronal function and thrashing frequency, we assessed the ‘Δ body length’ of worms expressing ChR2 under the *unc-17* or *unc-47* promotor (e.g. elongation and contraction for GABAergic and cholinergic stimulation respectively) as read-out for neuronal function after optogenetic stimulation with the OptoArm (1.6 mW/mm^2^) (Figure 9C). Subsequently the same worms were analyzed with the WF-NTP to assess their thrashing frequency (Figure 9D). Different worms from the same population were tested at D1, D4, D8 and D11 of adulthood. Worms expressing ChR2 under the *myo-3* promotor were included as a control for ‘Δ body length’. As previously described, we found that the postsynaptic muscle cells were relatively resistant to a functional decline in early life (Figure 9C, Liu et al., 2003). The cholinergic neurons, however, revealed an almost logistic decline in function. Strikingly, this decline was not observed in GABAergic neurons (Figure 9C). Then, we related the decline in thrashing frequency (Figure 9D) with the decline in neuronal functioning (Figure 9C) for each genotype (Figure 9E). While the functional decline of neurons in neither of the motoneuron populations correlated perfectly with the decline in thrashing frequency, the cholinergic neurons had a slope of almost 1, suggesting a strong relationship between the decline in locomotion and neuronal function (Figure 9E). Notice that the curve is slightly right-shifted (y = ax +b, b = ×0.267), which implies that, even though the *x, y* relationship is almost 1.0 (*x ∼ y*), a portion of the decline in motility cannot be explained by cholinergic deterioration alone. Therefore, our data suggests that cholinergic-dysfunction correlates best with the age-dependent movement deficits. At the same time, we show evidence for different ageing kinetics between the cholinergic and GABAergic motorneurons. GABAergic neurons appear more resilient to ageing, suggesting that not all *C. elegans* motoneurons age equally fast.

Since a delicate balance between GABAergic and cholinergic function is essential for movement, the disbalanced decay between the two-neuron population might contribute to a changed excitation-to-inhibition ratio to the muscle. As stated before, due to the wiring of the nervous system stimulation of cholinergic neurons normally causes a concurrent activation of GABAergic neurons, that results in typical coiling-behavior (McIntire et al., 1993; White et al., 1986; Schuske et al., 2004; Liewald et al., 2008; Figure 8), and provides a readout that captures both systems at once. Therefore, we hypothesized that a changed balance between the GABAergic and cholinergic system can translate into a changed light-induced coiling propensity. In order to tackle this question, we assessed the change in eccentricity (as a readout for coiling propensity) induced by illumination of worms expressing ChR2 in their cholinergic neurons at D4 and D8, just before and after the steep decline of the cholinergic output (Figure 9C). Although not significant, we observed a clear and consistent increase in ‘Δ eccentricity’ at D8 compared to D4 (Figure 9F). The decrease in eccentricity is almost twice as high (*Cohen’s d*: 1.99) at D8. In conclusions, we hypothesize that the observed increase in coiling propensity could be due to an inequal decline of cholinergic and GABAergic neurons and thus might provide a subtle hint for a changed excitation-to-inhibition ratio towards inhibition.

### Automation and additional features can further increase the ease of use and functionality of the OptoArm

All optogenetic data collected in this paper, was created with rather long photostimulation of at least 3 seconds. Those time-intervals can be relatively well established by manual operation of the OptoArm, e.g., using the ON-OFF switch. The manual OptoArm can definitely be valuable for acquiring consistent optogenetic data as is evidenced by all data that is described in this paper (Figure 7-9), but its consistency is highly operator-dependent. Moreover, it is practically impossible to generate the short-pulses or even high frequency trains required for certain experiments through manual operation (Liewald et al., 2008; Crawford et al., 2020). Therefore, we explored the use of a microcontroller to circumvent the use of a computer and to provide a low-cost solution to the aforementioned problems. The 700 mA externally dimmable BuckPuck DC driver that is used to power the LED, has a built-in 5V reference/output to directly power the logic circuitry of a *u*processor without the need for an additional power source (Figure 10A, Table 1). We used an Arduino uno as a microcontroller to control the output of the OptoArm, making it a stand-alone system. By connecting a pulse-width control pin (PWC, pin 9) to the CTRL pin of the driver, the Arduino cannot only control the duration of a light-pulse and the number of cycles, but can also adjust the light intensity, replacing the potentiometer (Figure 10B). These three parameters are easily adjustable by reprogramming of the Arduino software. However, adjusting Arduino scripts requires both knowledge of the programming language and a computer. Since we have tried to keep the OptoArm a standalone set-up that can be easily operated without specialistic knowledge and expensive equipment, we propose a more elegant solution. We decided to use an LCD-shield on top of the Arduino to adjust different parameters in live-modus without the intervention of a computer (Figure 10C). Then, we created an Arduino-script that allows users to use the LCD screen and connected buttons to select different options and menus (Supplementary software 1, Table 3, box 2). The software provides 3 programs through which the users can navigate for different experimental purposes: a testing program, a single run menu and a series pulse menu (Figure 10D,E, box 2). When using the automated OptoArm, one has to build the circuitry depicted in Figure 10A and upload the provided software to the Arduino only once (connect the CTRL pin of the LED driver to pin 9 of the Arduino to ensure compatibility between the software and the circuit). The used LCD-shield is relatively expensive when compared to the other components of the system, but it offers a user-friendly solution (Table 3, box 2). Nevertheless, other LCD screens can be used without a problem, as long as the software is adjusted as well.

**TABLE 3:** Arduino controls and button functions.

| **Display screen** | **Controls** | **Function** | **Fig. 10** |
| --- | --- | --- | --- |
| **Main** | UP/DOWN | Scroll through program options | D: I,III,V |
|  | SELECT | Confirm selection |  |
| **Program 1 settings** | UP/DOWN | Set intensity | D: II |
|  | LEFT/RIGHT | Switch between ON/OFF/MAIN |  |
|  | SELECT | Confirm return to main menu |  |
| **Program 2/3 settings** | LEFT/RIGHT | Set parameter value | D: IV, VI |
|  | SELECT | Confirm settings, skip to next display |  |
| **Program 3 settings** | UP/DOWN | Switch between parameters | D: VI |
| **Start run** | LEFT/RIGHT | Switch between yes/no | E: I |
|  | SELECT | Confirm selection |  |
| **Finish** | LEFT/RIGHT | Switch between rerun/return to main | E: II |
|  | SELECT | Confirm selection |  |

**Figure 10:**
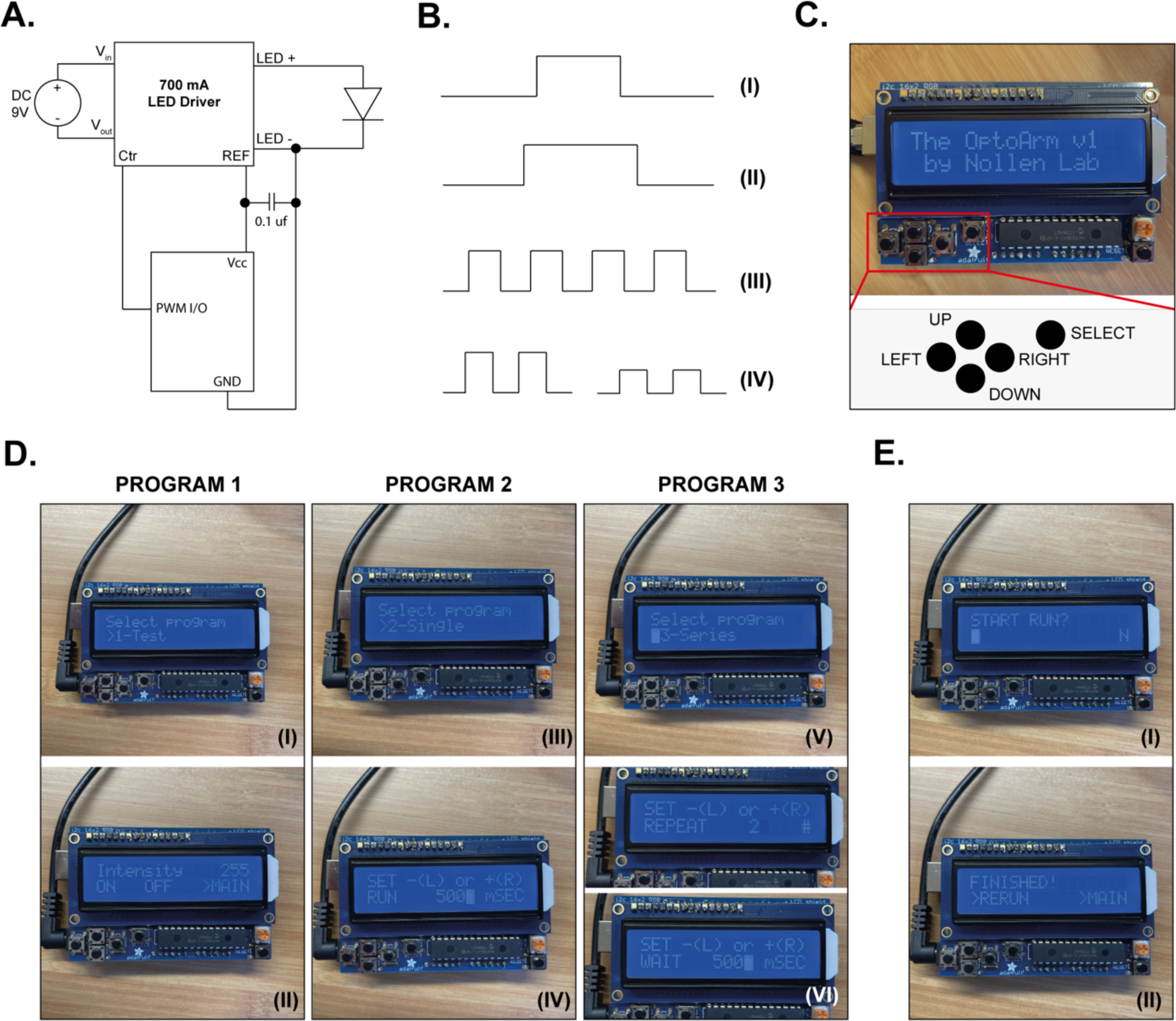
Automation of the OptoArm allows fine-adjustment of light pulse-width and intensity. **A)** The electronic circuitry of the OptoArm under the control of a micro-controller (an Arduino uno in this paper). **B)** By using a pulse-width modulator output pin of Arduino the pulse-width (I, II), time-intervals (III) and intensity (IV) op the OptoArm can be easily regulated. **C)** By using an LCD-shield on top of an Arduino uno, all parameters can be adjusted in live-modus without the intervention of a computer. There are 5 buttons that can be used to go UP and DOWN in the different programs to select parameters and to adjust them by pressing LEFT or RIGHT. All changes can be confirmed with the SELECT button (see Table 3). **D)** The different programs of the automated OptoArm. Program 1 (I) allows testing the system and manually turning the LED ON and OFF (II). Here, the intensity of the system can be changed as well. Program 2 (III) can be used to give single-timed pulses of light (IV). Program 3 (V) can be used to give trains of single-time pulses. One can adjust both the pulse-time (IV), the waiting time and the number of cycles (IV). **E)** The software provides a clear overview of the steps that are performed. When a run is started and then finishes, the user can use the option ‘Rerun’ to execute the same program without having to adjust the parameters again.

It has been underlined now several times that the OptoArm provides the user with an adaptable system for optogenetic experiments. The ease of automating the OptoArm strengthens that statement, but there is more to add. First of all, the system uses a royal-blue LED because ChR2 requires a wavelength in the blue spectrum. There are however many more, inhibitory light-gated ion pumps or outward-directed proton pumps that can be used for optogenetic purposes (Ihara et al., 1999; Waschuk et al., 2005; Zhang et al., 2007; Chow et al., 2010; Husson et al., 2012a). The yellow-green light-sensitive archaerhodopsin-3 (Arch) (Ihara et al., 1999), halorhodopsin from Natronomonas pharaonic (NpHR) (Zhang et al., 2007; Husson et al., 2012a) and rhodopsin from Leptosphaeria maculans (Mac) (Chow et al., 2010; Waschuk et al., 2005), are a few examples of the light-sensitive actuators that are widely used. NpHR, Arch and Mac are activated by wavelengths of 580-600 nm, 568 mm and 540 nm respectively (Husson et al., 2012a). By substituting the Royal Blue (448 nm) LED with a lime variant (567 nm), the OptoArm can also be used to photoactivate those inhibitory rhodopsins (Table 1). If one would be interested in switching between colors, we highly recommend to either built two arms, or to wire the OptoArm in such a way that only the heatsink and the attached LED can be disconnected and exchanged. Finally, while the OptoArm was specifically developed to work with *C. elegans*, it can also be used for experiments with *Drosophila melanogaster*. At present, required intensity for optogenetic stimulation of *D. melanogaster* is nowadays a few orders of magnitudes smaller. Commonly used intensities range from 50 *u*W/mm^2^ to 0.3 mW/mm^2^, although much higher intensities (<6.5 mW/mm2) are also used (Pulver et al., 2009; Dawydow et al., 2014; Meloni et al., 2020). To gain such small intensities with the OptoArm, one can use the systems intensity control, omit the use of lenses and/or increase the working-distance. In addition, with such a high sensitivity to light the effect of the backlight of recording systems (e.g. microscope and tracker lights) should be investigated as well.

## Discussion

While pharmacological tools and electrophysiology provide an unprecedented way to analyze synaptic function in *C. elegans,* optogenetic tools have revolutionized the way we can study neuronal circuits in a more non-invasive way (Husson et al., 2013). The expression of light-sensitive ion channels and pumps under tissue or cell-specific promotors, allow researchers to remotely modify and manipulate the behavior of excitable cells (Nagel et al., 2003; 2005; Li et al., 2005; Schroll et al., 2006; Bi et al., 2006; Ishizuka et al., 2006; Zhang et al., 2007; Fang-Yen et al., 2015; Husson et al., 2013). Consequently, the ability to tweak synaptic activity and concurrently observe its effect on behavioral readouts, makes it possible to dissect the neuronal circuits underlying animal behavior. While many systems are available for optogenetic purposes (Weissenberger et al., 2011; Leifer et al., 2011; Stirman et al., 2010; 2011; Pulver et al., 2011; Husson et al., 2012b; Kawazoe et al., 2013; Qiu et al., 2015; Rabinowitch et al., 2016b; 2016a; Gengyo-Ando et al., 2017; Pokala et al., 2018, Crawford et al., 2020) the associated costs, the required manual adaptations, compromises for usability and low adaptability provide a major bottleneck in the selection of the system. Here, we argue that optogenetics should be made accessible to all researchers regardless of limited financial resources or advanced technical expertise. Therefore, we designed a low cost optogenetics system that provides high quality and consistent experiments, while also being easy and flexible in useability.

We have developed the OptoArm as an inexpensive and simple to setup system for the optogenetic stimulation of *C. elegans*. It provides the user with adjustable light-intensity and lightning profiles via the incorporation of interchangeable lenses. The ability to change lenses allows optogenetic manipulation and analysis of the resulting behavior in both single worms and entire worm populations. Therefore, we technically validated the system in different setups, ensuring compatibility with both standard microscopes and existing worm-trackers like the WF-NTP. The compactness of the system makes it easy to incorporate in a variety of experimental setups, as the only requirement is a power outlet and minimal space. In addition, the OptoArm can be easily automated by the inclusion of a microcontroller, thereby allowing users to perform more complex optogenetic experiments involving high-frequency trains of light-pulses. By offering both the option to build a manual and a fully automated system, researchers are able to pick the set-up that fits their budget and requirements best. The cheapest version of the OptoArm only costs $65 dollar, while the variant with an included stand will take about $ 90 dollars in total. The fully automated system, including stand, is set to only ∼ $128 dollar, thereby outperforming most other optogenetic devises available. In addition, the capabilities of the system can be expanded by low-cost additional modules as the need presents itself, making it an investment for the long-term as well.

Considering the fact that overall costs are often a key determinant in adopting a new device, the OptoArm circumvents a major impediment. Next to its flexibility and adaptability, the OptoArm also bypasses the need of extra adjustments that are often required for low-cost systems, like aluminum foil or closed boxes, which are often incompatible with live imaging systems. Therefore, the OptoArm provides a way to illuminate and record worms simultaneously. Nevertheless, there are also some disadvantages when comparing the OptoArm with more expensive optogenetic systems (Leifer et al., 2011; Stirman et al., 2010; 2011; Weissenberger et al., 2011; Husson et al., 2012b; Qiu et al., 2015; Rabinowitch et al., 2016a;;Gengyo-Ando et al., 2017; Busack et al., 2020). First of all, the OptoArm does not allow to follow single worms in space with both temporal and chromatic precision. It does not allow illumination at a specified anatomical position only. Therefore, it is not possible to excite single neurons with the OptoArm. Nonetheless, this is only required for very specific, specialist applications. Secondly, our current configuration of the OptoArm provides an adaptable light intensity between ∼up till 1.8 mW/mm^2^ (with a fixed working-distance of 3.5 cm), but some experimental procedures require a higher intensity up till 5 mW/mm^2^ (Husson et al., 2013). However, it is possible to extend the system with a tri-LED that triples the power of the available light. In this way, with the appropriate thermal management and lenses, the light intensity of the OptoArm could be further increased. Thirdly, while it is possible to perform multi-worm illumination with the OptoArm, the spatial consistency is not ideal. For plates requiring illumination of a small surface area (e.g., 3-cm, 12-wells), the OptoArm can provide a coverage of sufficiently consistent light intensity. Yet, when larger areas are illuminated, the light-gradient and intensity drop towards the periphery becomes more of an issue. Lastly, the OptoArm cannot shift between wavelengths, as would be possible with a fluorescence microscope. Nevertheless, it is possible to change illumination color by incorporating different types of LEDs. In this way, the OptoArm could not only be used for photoactivation of ChR2, but also for Mac, NpHR or Arch (Husson et al., 2012a). Clearly, the modular nature of the OptoArm can be used to overcome some of the mentioned limitations.

While some disadvantages exist, we still demonstrated that the OptoArm can accurately reproduce previously established findings that were acquired with more expensive optogenetic systems. Furthermore, by using the OptoArm with the WF-NTP a variety of population characteristics can be acquired on both solid and liquid media, despite some spatial inconsistency in illumination. We also show that the OptoArm can be used to tackle novel questions by combining different optogenetic readouts. In fact, we used the OptoArm to study the age-related decline of specific motor neurons and the accompanying deterioration in locomotion. We found a striking correlation between the age-dependent decline in cholinergic output and the decline in thrashing capacity. However, the decrease in cholinergic output, as evidenced by a change in body length upon optogenetic stimulation, did not completely explain the drop in movement capacity during ageing. Neither did the decrease in GABAergic output or muscle function. In fact, we observed that the muscular and GABAergic system appeared to be more resilient to age-related deterioration. Indeed, the age-related decrease in GABAergic output was much smaller than that of the cholinergic branch. It has been previously shown that the overall function of a neuromuscular unit decreases early in life due to neuronal deficits rather than of muscle dysfunction (Liu et al., 2013). However, to our knowledge, the striking differences in ageing-kinetics between GABAergic and cholinergic neurons has not been described before.

The GABAergic and cholinergic system are inextricably linked and regulate the fine balance of excitation-to-inhibition stimuli to the muscle, ensuring coordinated movement. (White et al., 1986; McIntire et al., 1993). If GABAergic neurons are indeed more resilient to ageing, while cholinergic neurons gradually lose their functional capacity, the balance of the excitation-to-inhibition ratio for muscle cells might also change. Intriguingly, given the underlying interaction between GABAergic and cholinergic neurons that shapes the coiling phenotype (Liewald et al., 2008), the observed light-induced increase in coiling behavior at later ages might actually provide the first hint for an altered excitation-to-inhibition ratio. While the age-dependent decline in locomotion correlates well with a decrease in cholinergic signaling, it might be enhanced by the much slower decline in GABAergic function. To add another layer of complexity, a study by Liu et al. (2013) found evidence of postsynaptic sensitization, increasing the amplitude of both acetylcholine- and GABA-evoked muscle currents during early ageing and peaking at D9. The relationship between these postsynaptic adaptation mechanisms and presynaptic changes will further determine the delicate balance of the neuromuscular unit and the overall movement capacity. Clearly, further research to unravel the complex interplay that contributes to age-related decrease in movement capacity is required and should integrate all these components instead of focusing on a single compartment. Moreover, comparing GABAergic and cholinergic neurons molecularly might provide insight into the key players underlying the differences in resilience to ageing.

In conclusion, we have presented here a low-cost, easy-to-build and highly adaptable optogenetic device that allows researcher to perform novel optogenetic experiments with *C. elegans* and other small organisms. We validated the OptoArm technically and biologically in different set-ups for both single and multiple worm experiments. The flexible and adaptable nature ensures compatibility with different recording systems, allowing different readouts to be combined for more elaborate insight in biological systems. The OptoArm can be easily operated without expert knowledge and, due to its modular nature and cheap components, can be easily adapted as well. In the end, users have ultimate control over their budget and system configuration. Altogether, we show that the OptoArm is able to overcome some of the major obstacles preventing a more wide-spread implementation of optogenetics, including in teaching institutes.

## Materials and methods

### Strains and maintenance

Standard conditions were used for *C. elegans* propagation at 20 °C (Brenner, 1974). Animals were synchronized by hypochlorite bleaching, hatched overnight in M9 buffer and subsequently cultured on NGM agar plates seeded with OP50. For optogenetic experiments, transgenic worms were cultivated in the dark at 20 °C on NGM plates with or without 0.2 mM all-trans retinal (ATR) added to the OP50. A final concentration of 0.2 mM ATR was obtained by mixing 0.4 *u*l of a 100 mM ATR stock solution in ethanol (Sigma-Aldrich) with the 200 *u*L E. coli that was spread on each 6 cm plate. For ageing experiments, worms were transferred ∼72 hours after plating to NGM plates containing 5-Fluoro-2’deoxy-uridine (FUdR) to inhibit growth of offspring. For ageing experiments untill adulthood D12, worms were transferred every 3 days to fresh ATR+ and ATR-plates. The following genotypes were used: ZX388: zxIs3[*unc-47p::ChR2(H134R)::YFP + lin-15(+)]*V, ZX460: zxIs6[*unc-17p::ChR2(H134R)::YFP + lin-15(+)*]V, ZX463: *unc-47(e307)*III; *zxIs3*, ZX465: *unc-26(s1710)*IV; *zxIs3*, ZX511: *unc-26(s1710)*IV; *zxIs6*, ZX531: *unc-47(e307)*III; *zxIs6*. All strains were a kind gift of professor A. Gottschalk.

### The OptoArm

#### Required components

All the components of the Optoarm are by Quadica Developments, unless stated differently. For the electronic circuitry we refer to Figure 2A. The OptoArm consists of a Royal-Blue (448nm) Rebel LED on a SinkPAD-II 20 mm Star Base of 1.03 W (SP-01-V4), powered by a 700 mA externally dimmable BuckPuck DC driver – with leads (3023-D-E-700), and connected to a connecting wire with adjustable potentiometer of 5 kΩ (3021HEP). The LED is attached to a heatsink (50 mm Round x 44 mm high alpha heat sink – 5.2 °C/W; CN50-40B) with pre-cut thermal adhesive tape for 20 mm Star LED assemblies (LXT-S-12). The lens-holder was self-manufactured from aluminum with a Schaublin 125 lathe (*φ*21 mm, inner circle) and contains a small screw-hole at the side (M2) for tightening and loosening lenses and two grooves for the LED-wires (Figure 2B). The holder was attached to the heat sink with mounting tape for kathod 20 mm Round optic Holders (LT-06). Three different lenses (with adjacent integrated holder) were used: Khatod 10° 22mm circular beam optic (KEPL115406 by Khatod Optoelectronic), Fraen 21° 22 mm circular beam optic (FLP-M4-RE-HRF, by Fraen Corporation), Khatod 40° 22 mm circular beam optic (KEPL115440 by Khatod Optoelectronic). For the ON/OFF switched a latching pressure switch (≤3A, ≤ 250V; Velleman – R1821A-RD) was used. The whole system was mounted on a three-pronged clamp (*φ*10 mm, 241-7432, Usbeck Carl Fried). A cross clamp (105 BR 10, Comar) was placed on the rod of the clamp. The clamp provides a way to fixate a standard DC contra plug (≤4A, ≤25 V, Velleman; CD021) and to power the system with a DC-adapter. Due to the nature of a LED driver, the input voltage to the driver must always be higher than the total forward voltage drop of all connected and consuming components. In this case, the driver has a minimal input margin of 2.5 V and the LED consumes about 3.5 V, leading to a minimal input voltage of 6V. However, when working with a potentiometer it is highly recommended to use a minimum of 7V_DC_ (as evidenced by the spec sheet). We have been using a 9V_DC,_ 12V_DC_ and 15V_DC_ adapters (∼ 1.0 – 1.2 A) with good results. All components can also be found in table 1. Clearly, one can substitute the potentiometer, switch and DC connector for cheaper variants. The critical parts of the system include the LED, heat-sink, and steady 700mA driver together with the lenses.

#### Automated OptoArm

For the automated OptoArm, the aforementioned components are extended with an Arduino Uno (or any clone), a capacitor of 0.1 *u*f (Xicon Ceramic Disc Capacitors; Mouser electronics, 140-50U5-204M-RC) and an LCD Shield kit w/ 16×2 character display (Adafruit, ID:714), while the connecting wire with adjustable potentiometer of 5 kΩ can be omitted. The shield has to be assembled with some soldering-steps; a great tutorial can be found here: https://learn.adafruit.com/rgb-lcd-shield/assembly. See Figure 10A for the electronic circuitry. The accompanying Arduino code for the LCD-shield is freely available and can be downloaded from: Supplementary Software 1 (the software runs under the license of Attribution-NonCommercial-ShareAlike 4.0 International (CC BY-NC-SA 4.0). When using this software, connect pin D9 of the Arduino to CTRL pin of the LED driver.

#### Stands

For experiments with the WF-NTP, we used a stand that was built with the following components: stand rod (1000 mm, *φ*12 mm, 241-7153, Usbeck Carl Fried), double bosshead (241-7104, Usbeck Carl Fried), retort stand base, tripod (241-0245, Usbeck Carl Fried). For experiments with a microscope, we used a stand with the following components: basic carrier (20 RM 01, Comar), Pinion stage (30 XT 40), post-holder (45 BH 10), rod (203 RM 01), self-manufactured base.

### Microscope and WF-NTP setup

Microscope: a high-resolution microscopy system was used to image the worms, consisting of an Olympus SZ51 microscope coupled with an IphoneX via a Carson HookUpz 2.0. smartphone adapter. The UV-protection shield of a Leica MZ 16 FA was used to avoid saturation of the camera. In general, a 1.7x digital zoom (of the Iphone) was used in combination with a 15-30x magnification of the microscope, and videos were always acquired with a framerate of 30 fps. Actual working distances were determined by measuring the height from the surface perpendicular to the middle of the LED and using the trigonometric ratio for sine, eq. 9, to calculate the length of the hypotenuse side:

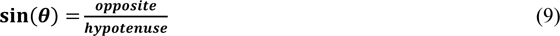

A working-distance of 3.5 cm is often adequate. WF-NTP: the used set-up was identical to the one described in Koopman et al., (2020), using a camera-to-sample distance of 140 mm. The used framerate was always 20 fps and the working-distance was set to ∼ 4.1 cm. In both set-ups, light intensity was always measured at 448 nm with a M16-130 USB power meter (Thorlabs, *φ* 9.5 mm) to verify the required conditions. Before each experiment we measured the light-intensity to ensure appropriate conditions, assuming a Gaussian light source (in the software).

### Behavioral experiments

For single-worm illumination, worms were freely moving on unseeded 3-cm NGM plates and kept in frame by manual location. For ‘short-term’ experiments, worms were exposed to a 3-3-3 lightening regime, in which light was OFF for 3 seconds, followed by light ON for 3 seconds and light OFF for another 3 seconds. Light intensity was always 1.6 mW/mm^2^ (10° lens) unless stated differently. For ‘long-term’ experiments, worms were exposed to a 1-50-1 lightening regime in which light was OFF for 1 seconds, followed by light ON for 50 seconds and light OFF for another second. For multiple-worm illumination, worms were collected in M9 buffer and plated on an empty 3-cm plate that was flooded with 1.5 mL M9. Worms were recorded with the WF-NTP set-up at a framerate of 20 fps and a light intensity of 1.0 mW/mm^2^ (21° lens) unless stated differently. We generated separate movies for light OFF (30 seconds) and light ON (30 seconds). For most experiments, as for the ageing-timeline, worms were first examined via single-worm illumination and subsequently transferred to M9 for multi-worm illumination.

### Image processing

Videos of multiple-worm illumination, acquired with the WF-NTP platform, were analyzed with accompanying software as described in Koopman et al., 2020. Videos that were acquired with a cell-phone (Iphone X) were transferred to a computer (.mov) and first converted to ImageJ compatible files (.avi) with ffmpeg. We used a terminal (cmd for Windows, terminal for MacOS) to navigate to the directory with the movies and quickly convert multiple movies. When opening a terminal, the current location will be visible (e.g., C:\Users\). When the movies are saved at a different disk, for example, D, one can use the command: C:\Users\Tracker>**d**: followed by pressing ‘Enter’ to switch directly to that disk. Paths can be removed with the command **cd\,** for example, C:\Users\Movies>**cd\** followed by pressing ‘Enter’. This will yield ‘C:\Users’. Adding paths is managed by typing cd **X**, for example, C:\>cd **Movies** followed by pressing ‘Enter’. This will yield ‘C:\**Movies**. When the appropriate directory is selected, the following command can be executed:

for i in *.MOV; do ffmpeg -i “$i” -pix_fmt nv12 -f avi -vcodec rawvideo “${i%.*}.avi”; done

This code converts all .MOV files in the directory to .AVI files at the same time. For the perimeter-approach, the converted movies were processed with ImageJ. Movies were first set to an 8-bit format (Image > Type > 8-bit) and then duplicated (Image > Duplicate) to ensure accessible images for all subsequent steps. Next, the movie contrast was enhanced by selecting 0.3% saturated pixels and normalized enhancement (Process > Enhance), followed by background subtraction with a rolling ball radius of 50 px (Process > Subtract Background). Then, we thresholded the images (Image > Adjust > Threshold) by the Huang method (same threshold for the wole stack) and applied the settings to all frames. Finally, we measured the perimeter of the worm per frame by setting the measurements (Analyze > Set measurements) to perimeter only and clicking on ‘analyze particles’ (Analyze > Analyze particles). One should set a lower-limit of the size of the particles to avoid noise being picked up. This number highly depends on the used magnification and should therefore be determined by an empirical approach. We recommend to record the first processing steps until subtracting the background (Plugins > Macros > Record) and created a macro (Create) that can be rerun for every movie. In this way, only the thresholding had to be performed manually. Measurements were normalized by the average pixel length of the perimeter measured in the first 30-60 frames (light OFF). For long-term experiments, a midline approach was used at specified intervals (0.5 seconds, 2 seconds, 10 seconds, 20 seconds, 30 seconds, 40 seconds, 50 seconds and 52 seconds). If the midline approach was used for short-term experiments, we measured the body-length 500 ms before light was ON and 1000 ms after light was turned ON. When the midline-approach is specifically mentioned, we measured the body length by drawing a midline from head to tail with a drawing-tablet (Wacom).

### Statistics and visualization

Statistical analyses were done in R and Graphpad Prism 8. The used statistical tests can be found in the different figures and are based on several criteria. In short, (log-)normality tests were always performed on the collected data to test for gaussian distributions (e.g. Shapiro-Wilk). Based on the distribution of the data the appropriate test was selected (parametric or non-parametric). When more than two groups were compared, multiple-testing correction was always applied. When inequality of variance was expected between two groups (based on the experimental design) student’s t-test were always performed with a Welsch’s correction (The Welch test must be specified as part of the experimental design, and not decided upon *a posterori;* Moser et al., 1992; Hayes et al., 2007). Post-hoc testing after a one- or two-way ANOVA was only performed when the initial test gave significant results. All experiments were replicated three times, unless stated differently. *: p ≤ 0.05, **: p ≤ 0.01, ***: p ≤ 0.001. All data was visualized with R (ggplot2) or Graphpad Prism 8 and color-adjusted in Adobe Illustrator. Integrated intensity plots were created with R, by applying a general LOESS model on the measured point to estimate/interpolate the intensities at the non-measured coordinates.

## Supporting information

Supplementary Software 1

## Acknowledgements

We thank Alexander Gottschalk for kindly sharing several strains and fruitful tips and trick on optogenetics with *C. elegans*. This project was funded by a BCN-Brain grant (to M.K.) and a grant of Stichting ALS Nederland (to E.A.A.N. and M.K.)

## Author contributions

L.J. and M.K. designed and built the OptoArm together. M.K. performed all technical and biological experiments, analyzed and visualized them. L.J. wrote the accompanying Arduino script. M.K. wrote the manuscript with input from L.J. and E.A.A.N.

## Conflict of Interests

The authors declare that they have no conflict of interest.

## Boxes

#### BOX 1: Calculating the required thermal properties of a heat sink

Using the following thermal model (eq. 1, Lin et al., 2011, Figure 2F) and calculating or estimating all the individual parameters (eq. 2-4) the required thermal properties of a heat sink can be calculated.

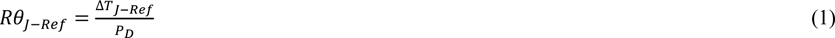

Where:

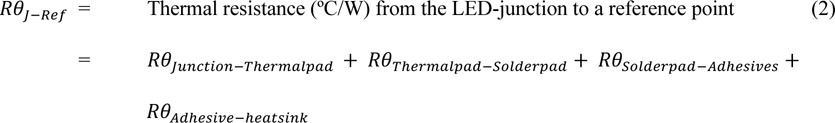

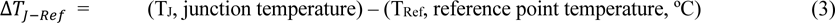

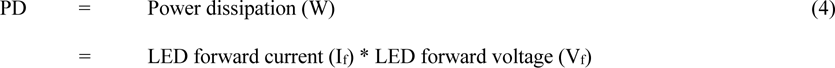

Typically, the maximal junction temperature can be found in the datasheet for the LED. For a Royal blue Luxeon Rebel LED on a SinkPAD-II, that is used in this paper, the listed temperature is 150 °C. Ideally, the operating temperature should be well below that limit, as even reaching this temperature for a fraction of time could influence the properties of the LED (Yang et al., 2018; Oh et al., 2019; Fu et al., 2018; Kim, et al., 2016). Therefore, we decided to set the maximum junction temperature to 100 °C. Next, the power dissipation of the same LED can be calculated by multiplying the forward voltage rating of the LED with the drive current in Amperes (eq. 4). Since we use a fixed 700 mA LED-driver and know the maximal voltage drop of the LED (3.5V), P_d_ is easily calculated (eq. 4). Finally, the maximal thermal resistance should be calculated by taking in the maximum working ambient temperature into account, which was determined to be 25 °C (20 °C room temperature + 5 °C margin, eq. 3). Importantly, note that our set-up is located in a climate-controlled room. If this is not the case, a wider margin for temperature fluctuations and higher maximum temperature based on local environmental conditions should be considered. Rewriting eq. 1 and calculating *Rθ* gives (eq. 5):

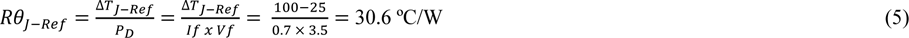

This means that the total thermal resistance of the system can be maximally 30.6 °C/W (eq. 2) when one aims for junction temperatures that are maximally 100 °C. In order to see what that means for the required heatsink, all the known thermal resistances (Figure 2F) can be collected from the datasheets of the LED (junction to thermal pad: 6.0 °C/W and thermal pad to solder pad: 0.7 °C/W) and used adhesives (4.5 °C/W). Rewriting eq 2. gives (eq. 6):

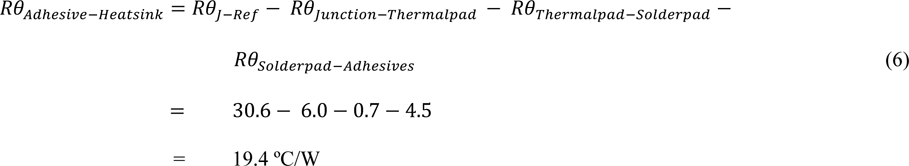

Theoretically, a heatsink in this set-up can have a maximal thermal resistance of 19.4 °C/W and any heatsink with lower resistance than that can be used. We used a finned heatsink with thermal resistance of 5.2 °C/W to keep the set-up compact and still achieve sufficient cooling. Next, we continued with an empirical approach to verify whether appropriate heat dissipation could be obtained with this set-up. Therefore, we measured the actual junction temperature and the voltage drop of the LED mounted to the OptoArm (Figure 2D). With a Fluke TM80 module and associated test probe, we measured the temperature at the specified test-location (Figure 2F) of the LED for about 1 minute until the temperature stabilized. At the same temperature, we also determined the actual voltage drop of the LED. We measured a temperature of 55.68 ± 0.76 °C (*n = 10*) and a forward drop of 2.972 ± 0.004 V (*n = 5*). By using eq. 7, the actual junction temperature can be derived:

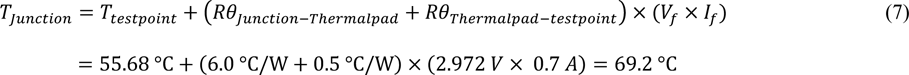

With an estimated junction temperature of 69.2 °C, the system fulfills our criteria for a system with appropriate thermal cooling. This is further evidence by the temperature measured after 10 minutes of constant illumination at the maximal capacity: 55.81 ± 1.09 °C (=junction temperature of 69.3 °C, *n = 10*).

#### BOX 2: The different programs within the OptoArm software

*Program 1:* The testing menu. This menu is used to set-up your experiment. The menu allows the user to select the intensity of the LED by changing the analog scale: 0-255 (buttons up and down) and to turn the LED ON at that intensity. This menu is ideal for adjusting the working-distance and incidence angle to gain the appropriate intensity before starting an actual experiment. Importantly, as long as the Arduino is powered, the set intensity will be saved and the standard for the other menu-options (Figure 10D – I/II, Table 2).

*Program 2:* The single run menu. In this menu the user can select the time of the light-pulse (Figure 10D – III/IV). The buttons are time-sensitive: the longer one pushes, the steeper the steps. There is standard a 1 second waiting step before the pulse when the run is started. When the run is completed, the users get the opportunity to rerun the assay, without setting the parameters again (Figure 10E, Table 2).

*Program 3:* The series pulse menu. In this menu the user cannot only select the time for the light-pulse, but also the waiting time (between two light-pulses) and the number of cycle-repeats (Figure 10D – IV-VI). By going up and down in the menu, the different parameters can be adjusted (Table 2). There is again a 1 second waiting step before the first pulse when the run is started. When the run is completed, the users get the opportunity to rerun the assay, without setting the parameters again (Figure 10E).

## Notes

### Competing Interest Statement

The authors have declared no competing interest.

### Summary of Updates

Incorrect names in the acknowledgement section.

