## Supplementary Software 1 for "Optogenetic manipulation of individual or whole population *Caenorhabditis elegans* worms with an under hundred-dollar tool: the OptoArm": OptoArm_controller_v1.ino

```
/*
  Set up basic lcd menus and allows setting of options to control LED for optogenetics
 :Author: Leen Janssen
 :
 :Date: 5/02/2021
 :Revision: version 1.0
 :License: Attribution-NonCommercial-ShareAlike 4.0 International (CC BY-NC-SA 4.0)
*/

// include the library code:
//I2C
#include <Wire.h>
//Lcd shield library and I2c Port Expander
#include <Adafruit_RGBLCDShield.h>
#include <utility/Adafruit_MCP23017.h>


// The LCD shield uses the I2C SCL and SDA pins. On classic Arduinos
// this is Analog 4 and 5 so you can't use those for analogRead() anymore
// However, you can connect other I2C sensors to the I2C bus and share
// the I2C bus. (LCD Shield default 0x20)
Adafruit_RGBLCDShield lcd = Adafruit_RGBLCDShield();

// These #defines make it easy to set the backlight color
#define OFF 0x0
#define ON 0x1

//class to handle button input for various menus
class Manager {
  public:
    //keeps track if buttons were pressed sice last check
    bool changed = false;
    //menustate 0= startmenu, 1 = set intensity, 2= set parameters, 3= start run, 4=running,5=finished
    int menuState = 0;
    //varies per menu, usually: -1 = skip to next menu, 1 = change value in menu, =cursorY to select menu item
    int optionChanged = 0;
    // to position cursor on lcd screen
    int cursorX = 0;
    int cursorY = 1;
    //to set parameters for custom program
    int sRun = 500;
    int sWait = 500;
    int steps = 0;
    int brightness = 255;
    int increment = 1;

    // Constructor - creates a View object
    // and initializes the member variables and state
  public:
    Manager(int menu) {
      menuState = menu;
    }

    //Functions for each button press in the various menu screens (=menuState)
    //may need to be revised changes are made to display text
    void Select() {
      switch (menuState) {
        case 0 :
          optionChanged = cursorY;
          break;
        case 1 :
          if (cursorX == 11)
          {
            optionChanged = -1;
          }
          break;
        case 2 :
          optionChanged = -1;
          break;
        case 3 :
          if (cursorX == 0)
          {
            optionChanged = 1;
          }
          else
          {
            optionChanged = -1;
          }
          break;
        case 5 :
          if (cursorX == 0)
          {
            optionChanged = 1;
          }
          else
          {
            optionChanged = -1;
          }
          break;
        default :
          optionChanged = 0;
      }
      changed = true;
    }

    void Down(int p) {
      if (p > 3) {
        increment = 5;
      }
      else {
        increment = 1;
      }
      switch (menuState) {
        case 0 :
          if (cursorY == 3 )
          {
            cursorY = 1;
          }
          else
          {
            cursorY = cursorY + 1 ;
          }
          break;
        case 1 :
          brightness = brightness - increment;
          if (brightness <= 0)
          {
            brightness = 255;
          }
          optionChanged = 1;
          break;
        case 2 :
          if (steps > 0) {
            if (cursorY == 2)
            {
              cursorY = 0;
            }
            else
            {
              cursorY = cursorY + 1;
            }
            optionChanged = 1;
          }
          break;
        default :
          cursorY = 0;

      }
      changed = true;
    }

    void Up(int p) {
      if (p > 3) {
        increment = 5;
      }
      else {
        increment = 1;
      }
      switch (menuState) {
        case 0 :
          if (cursorY == 1 )
          {
            cursorY = 3;
          }
          else
          {
            cursorY = cursorY - 1;
          }
          break;
        case 1 :
          brightness = brightness + increment;
          if (brightness >= 255)
          {
            brightness = 0;
          }
          optionChanged = 1;
          break;
        case 2 :
          if (steps > 0) {
            if (cursorY == 0)
            {
              cursorY = 2;
            }
            else
            {
              cursorY = cursorY - 1;
            }
            optionChanged = 1;
          }
        default :
          cursorY = 0;
      }
      changed = true;
    }

    void Left(int p) {
      if (p > 12) {
        increment = 100;
      }
      else if (p > 3) {
        increment = 10;
      }
      else {
        increment = 1;
      }
      switch (menuState) {
        case 1 :
          if (cursorX == 0)
          {
            cursorX = 11;
          }
          else if (cursorX == 11)
          {
            cursorX = 5;
          }
          else
          {
            cursorX = 0;
          }
          optionChanged = 1;
          break;
        case 2 :
          cursorX = 8;
          switch (cursorY) {
            case 0:
              sRun = sRun - increment;
              if (sRun <= 0)
              {
                sRun = 10000;
              }
              break;
            case 1:
              sWait = sWait - increment;
              if (sWait <= 0)
              {
                sWait = 10000;
              }
              break;
            case 2:
              steps = steps - increment;
              if (steps < 0)
              {
                steps = 0;
              }
              break;
          }

          break;
        case 3:
          if (cursorX == 0)
          {
            cursorX = 15;
          }
          else
          {
            cursorX = 0;
          }
          break;
        case 5 :
          if (cursorX == 0)
          {
            cursorX = 11;
          }
          else
          {
            cursorX = 0;
          }
          break;
        default :
          cursorX = 0;

      }
      changed = true;
    }

    void Right(int p) {
      if (p > 12) {
        increment = 100;
      }
      else if (p > 3) {
        increment = 10;
      }
      else {
        increment = 1;
      }
      switch (menuState) {
        case 1 :
          if (cursorX == 0)
          {
            cursorX = 5;
          }
          else if (cursorX == 5)
          {
            cursorX = 11;
          }
          else
          {
            cursorX = 0;
          }
          optionChanged = 1;
          break;
        case 2 :
          cursorX = 8;
          switch (cursorY) {
            case 0:
              sRun = sRun + increment;
              if (sRun > 10000)
              {
                sRun = 0;
              }
              break;
            case 1:
              sWait = sWait + increment;
              if (sWait > 10000)
              {
                sWait = 0;
              }
              break;
            case 2:
              steps = steps + increment;
              if (steps > 100)
              {
                steps = 100;
              }
              break;
          }
        case 3:
          if (cursorX == 0)
          {
            cursorX = 15;
          }
          else
          {
            cursorX = 0;
          }
          break;
        case 5 :
          if (cursorX == 0)
          {
            cursorX = 11;
          }
          else
          {
            cursorX = 0;
          }
          break;
        default :
          cursorX = 0;
      }
      changed = true;
    }

    //resets manager parameters to begin values
    void Reset() {
      menuState = 0;
      cursorY = 1;
      cursorX = 0;
      optionChanged = 0;
      changed = false;
      sRun = 500;
      sWait = 500;
      steps = 0;
    }
};


//Initialize class to handle button input
Manager dispM(0);
//set string arrays for menu items
char* mainItems[3] = {">1-Test         ", ">2-Single      ", ">3-Series       "};
char* subMenuL1[5] = {"Intensity       ", "SET -(L) or +(R) or", "START RUN?      ", "RUNNING...      ", "FINISHED!        "};
char* subMenuL2[4] = {"ON   OFF   >MAIN", "RUN         mSEC", "Y              N", ">RERUN     >MAIN"};
char* ParameterMenuL2[3] = {"RUN         mSEC", "WAIT        mSEC", "REPEAT         #"};
//set led pin
int led_pin = 9;
//keep track of duration button press
int up = 0;
int down = 0;
int left = 0;
int right = 0;


void setup() {
  // Debugging output
  Serial.begin(9600);
  // set up the LCD's number of columns and rows:
  lcd.begin(16, 2);
  analogWrite(led_pin, 255);

  //start up lcd screen
  lcd.print("The OptoArm v1");
  lcd.setCursor(0, 1);
  lcd.print(" by Nollen Lab");
  lcd.setBacklight(ON);
  delay(2000);
  lcd.clear();
  lcd.setCursor(0, 0);
  lcd.print("Select program");
  lcd.setCursor(dispM.cursorX, dispM.cursorY);
  lcd.print(mainItems[0]);
  lcd.blink();
  lcd.setCursor(dispM.cursorX, dispM.cursorY);

}
//uint8_t i=0;
void loop() {
  //check for button presses
  // delay of 200ms to debounce button press
  uint8_t buttons = lcd.readButtons();

  if (buttons) {
    if (buttons & BUTTON_UP) {
      dispM.Up(up);
      delay(200);
      up = up + 1;
      down = 0;
      left = 0;
      right = 0;
    }
    if (buttons & BUTTON_DOWN) {
      dispM.Down(down);
      delay(200);
      down = down + 1;
      up = 0;
      left = 0;
      right = 0;
    }
    if (buttons & BUTTON_LEFT) {
      dispM.Left(left);
      delay(200);
      left = left + 1;
      up = 0;
      down = 0;
      right = 0;
    }
    if (buttons & BUTTON_RIGHT) {
      dispM.Right(right);
      delay(200);
      right = right + 1;
      up = 0;
      down = 0;
      left = 0;
    }
    if (buttons & BUTTON_SELECT) {
      dispM.Select();
      delay(200);
      up = 0;
      down = 0;
      left = 0;
      right = 0;
    }
  }
  else {
    up = 0;
    down = 0;
    left = 0;
    right = 0;
  }

  if (dispM.menuState == 4) {
    dispM.changed = true;
  }

  //change lcd according to user input
  if (dispM.changed) {
    switch (dispM.menuState) {
      case 0 :
        if (dispM.optionChanged == 0) {
          lcd.setCursor(0, 1);
          lcd.print(mainItems[dispM.cursorY - 1]);
          lcd.setCursor(0, 1);
        }
        else {
          dispM.menuState = dispM.cursorY;
          if (dispM.menuState == 1) {
            lcd.clear();
            lcd.setCursor(0, 0);
            lcd.print(subMenuL1[dispM.cursorY - 1]);
            lcd.setCursor(0, 1);
            lcd.print(subMenuL2[dispM.cursorY - 1]);
            lcd.setCursor(13, 0);
            lcd.print(dispM.brightness);
            lcd.setCursor(5, 1);
            dispM.cursorY = 1;
            dispM.cursorX = 5;
          }
          else {
            dispM.cursorY = 2;
            lcd.clear();
            lcd.setCursor(0, 0);
            lcd.print(subMenuL1[dispM.cursorY - 1]);
            lcd.setCursor(0, 1);
            lcd.print(subMenuL2[dispM.cursorY - 1]);
            lcd.setCursor(7, 1);
            lcd.print(dispM.sRun);
            if (dispM.menuState == 3) {
              dispM.steps = 2;
            }
            else {
              dispM.steps = 0;
            }
            dispM.cursorY = 0;
            dispM.cursorX = 7;
            dispM.menuState = 2;
          }
          dispM.optionChanged = 0;
        }
        break;
      case 1 :
        if (dispM.optionChanged == -1) {
          /*back to main menu*/
          dispM.Reset();
          lcd.clear();
          lcd.setCursor(0, 0);
          lcd.print("Select program");
          lcd.setCursor(dispM.cursorX, dispM.cursorY);
          lcd.print(mainItems[0]);
          lcd.blink();
          lcd.setCursor(dispM.cursorX, dispM.cursorY);
        }
        else if (dispM.optionChanged == 1) {
          lcd.setCursor(13, 0);
          lcd.print("   ");
          lcd.setCursor(13, 0);
          lcd.print(dispM.brightness);
          switch (dispM.cursorX) {
            case 0:
              lcd.setCursor(0, 1);
              analogWrite(led_pin, 255-dispM.brightness);
              break;
            case 5:
              lcd.setCursor(5, 1);
              analogWrite(led_pin, 255);
              break;
            case 11:
              lcd.setCursor(11, 1);
              analogWrite(led_pin, 255);
              break;
          }
        }
        dispM.optionChanged = 0;
        break;
      case 2 :
        if (dispM.optionChanged == -1) {
          dispM.menuState = 3;
          lcd.clear();
          lcd.setCursor(0, 0);
          lcd.print(subMenuL1[dispM.menuState - 1]);
          lcd.setCursor(0, 1);
          lcd.print(subMenuL2[dispM.menuState - 1]);
          lcd.setCursor(0, 1);
          dispM.cursorY = 1;
          dispM.cursorX = 0;
        }
        else
        {
          if (dispM.steps == 0) {
            lcd.setCursor(0, 1);
            lcd.print(ParameterMenuL2[0]);
            lcd.setCursor(7, 1);
            lcd.print("   ");
            lcd.setCursor(7, 1);
            lcd.print(dispM.sRun);
          }
          else
          {
            lcd.setCursor(0, 1);
            lcd.print(ParameterMenuL2[dispM.cursorY]);
            lcd.setCursor(7, 1);
            lcd.print("    ");
            switch (dispM.cursorY) {
              case 0:
                lcd.setCursor(7, 1);
                lcd.print(dispM.sRun);
                break;
              case 1:
                lcd.setCursor(7, 1);
                lcd.print(dispM.sWait);
                break;
              case 2:
                lcd.setCursor(10, 1);
                lcd.print(dispM.steps);
                break;
            }
          }
        }
        dispM.optionChanged = 0;
        break;
      case 3 :
        if (dispM.optionChanged == -1) {
          /*back to main menu*/
          dispM.Reset();
          lcd.clear();
          lcd.setCursor(0, 0);
          lcd.print("Select program");
          lcd.setCursor(dispM.cursorX, dispM.cursorY);
          lcd.print(mainItems[0]);
          lcd.blink();
          lcd.setCursor(dispM.cursorX, dispM.cursorY);
        }
        else if (dispM.optionChanged == 1)
        {
          dispM.menuState = 4;
        }
        else {
          lcd.setCursor(dispM.cursorX, 1);
        }
        dispM.optionChanged = 0;
        break;
      case 4 :
        lcd.clear();
        //include 1 sec wait before start
        lcd.setCursor(11, 1);
        lcd.print("WAIT");
        delay(1000);
        lcd.clear();
        lcd.setCursor(0, 0);
        lcd.print(subMenuL1[dispM.menuState - 1]);
        if (dispM.steps == 0) {
          analogWrite(led_pin, 255-dispM.brightness);
          delay(dispM.sRun);
          analogWrite(led_pin, 255);
        }
        else {
          for (int s = 1; s <= dispM.steps; s++) {
            lcd.setCursor(0, 1);
            lcd.print("Step ");
            lcd.print(s);
            lcd.print("       ON");
            analogWrite(led_pin, 255-dispM.brightness);
            delay(dispM.sRun);
            analogWrite(led_pin, 255);
            lcd.setCursor(11, 1);
            lcd.print("WAIT");
            delay(dispM.sWait);
          }
        }
        lcd.clear();
        dispM.menuState = 5;
        lcd.setCursor(0, 0);
        lcd.print(subMenuL1[dispM.menuState - 1]);
        lcd.setCursor(0, 1);
        lcd.print(subMenuL2[dispM.menuState - 2]);
        lcd.setCursor(0, 1);
        break;
      case 5 :
        if (dispM.optionChanged == -1) {
          /*back to main menu*/
          dispM.Reset();
          lcd.clear();
          lcd.setCursor(0, 0);
          lcd.print("Select program");
          lcd.setCursor(dispM.cursorX, dispM.cursorY);
          lcd.print(mainItems[0]);
          lcd.blink();
          lcd.setCursor(dispM.cursorX, dispM.cursorY);
        }
        else if (dispM.optionChanged == 1) {
          dispM.menuState = 4;
        }
        else {
          lcd.setCursor(dispM.cursorX, 1);
        }
        dispM.optionChanged = 0;
        break;
      default :
        ;
    }
    dispM.changed = false;
  }
}
```
