## Supplementary Software 1 for "Optogenetic manipulation of individual or whole population *Caenorhabditis elegans* worms with an under hundred-dollar tool: the OptoArm": Readme.txt

:Author: Leen Janssen
:
:Date: 5/02/2021
:Revision: version 1.0
:License: Attribution-NonCommercial-ShareAlike 4.0 International (CC BY-NC-SA 4.0)

= Project: OptoArm 

This code was designed to allow the stand-alone operation of an optogentic set-up: the OptoArm. 
Allowing the user to select an appropriate program and set individual parameters, such as intensity, duration and number of light cycles, using an lcd display module for input . 


== Step 1 : Assemble the circuit

See paper by Koopman et al. for the circuit diagram to include Arduino in OptoArm set-up.
The code uses PWM pin 9 for connecting the LED. PWM is necessary to allow intensity regulation.
Other PWM pins can be used if preferred. Simply connect LED to other different PWM pin and change pin number on line 361 of the code. 

**Try testing connections and components of the construction to rule out faulty construction if issues occur later on.

== Step 2: Import the code
Import the sketch into an Arduino compatible editor, like the Arduino web editor.

== Step 2bis (optional): Edit the code
Depending on your systems requirements and personal preferences you can edit the code.
Please take note that changes to the code can cause the code to malfunction. Sometimes in other, perhaps unexpected places.
For instance, changes to the display text may cause issues accurate display of the cursor and menu items.

== Step 3: Connect Arduino to PC
Connect Arduino using USB-cable

== Step 4: Load the code

Upload the code on to your board

== Step 5: Test set-up

For a more detailed explanation of step 2-4 please go to
https://www.arduino.cc/en/Guide/ArduinoUno

=== Contributing
To contribute to this project please contact Leen Janssen <>
